## Supplementary Data for "Simultaneous inhibition of DNA-PK and Polϴ improves integration efficiency and precision of genome editing"

**Supplementary Figure S1.**

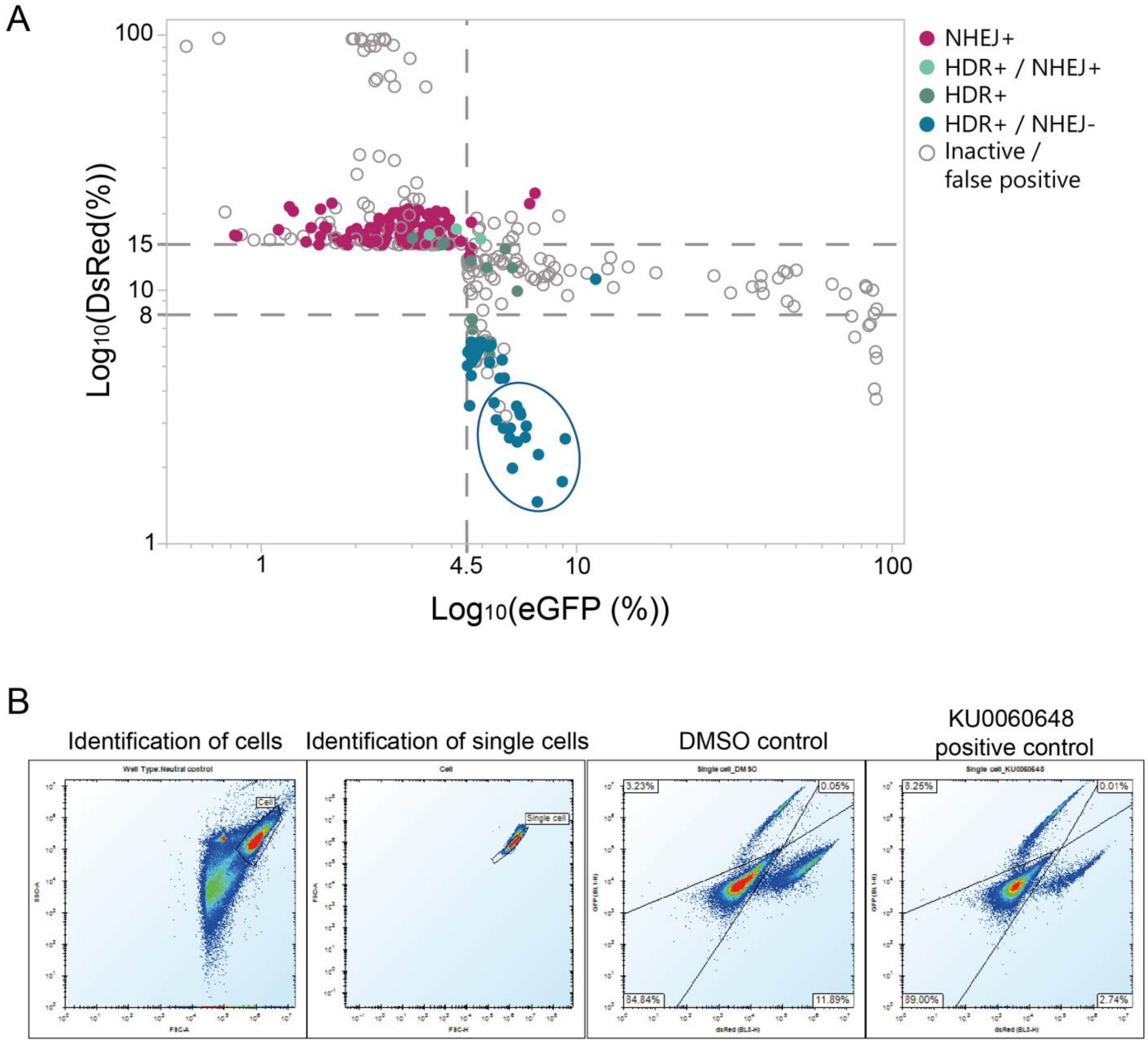

**Supplementary Figure S1 related to Figure 1: Validation of primary high-throughput screening results by a hit confirmation screen.** (**A**) Dots represent revised categories of the primary screen values by the outcome of the hit confirmation screen (n =1). (**B**) Representative flow cytometry plots of DMSO and KU0060648 treated cells. Gating strategy for cells based on forward scatter (FSC) and side scatter (SSC), subsequently gated for singlets with FSC height (FSC-H) and FSC-area (FSC-A). Followed by eGFP and dsRed quantification in the single cell population.

**Supplementary Figure S2.**

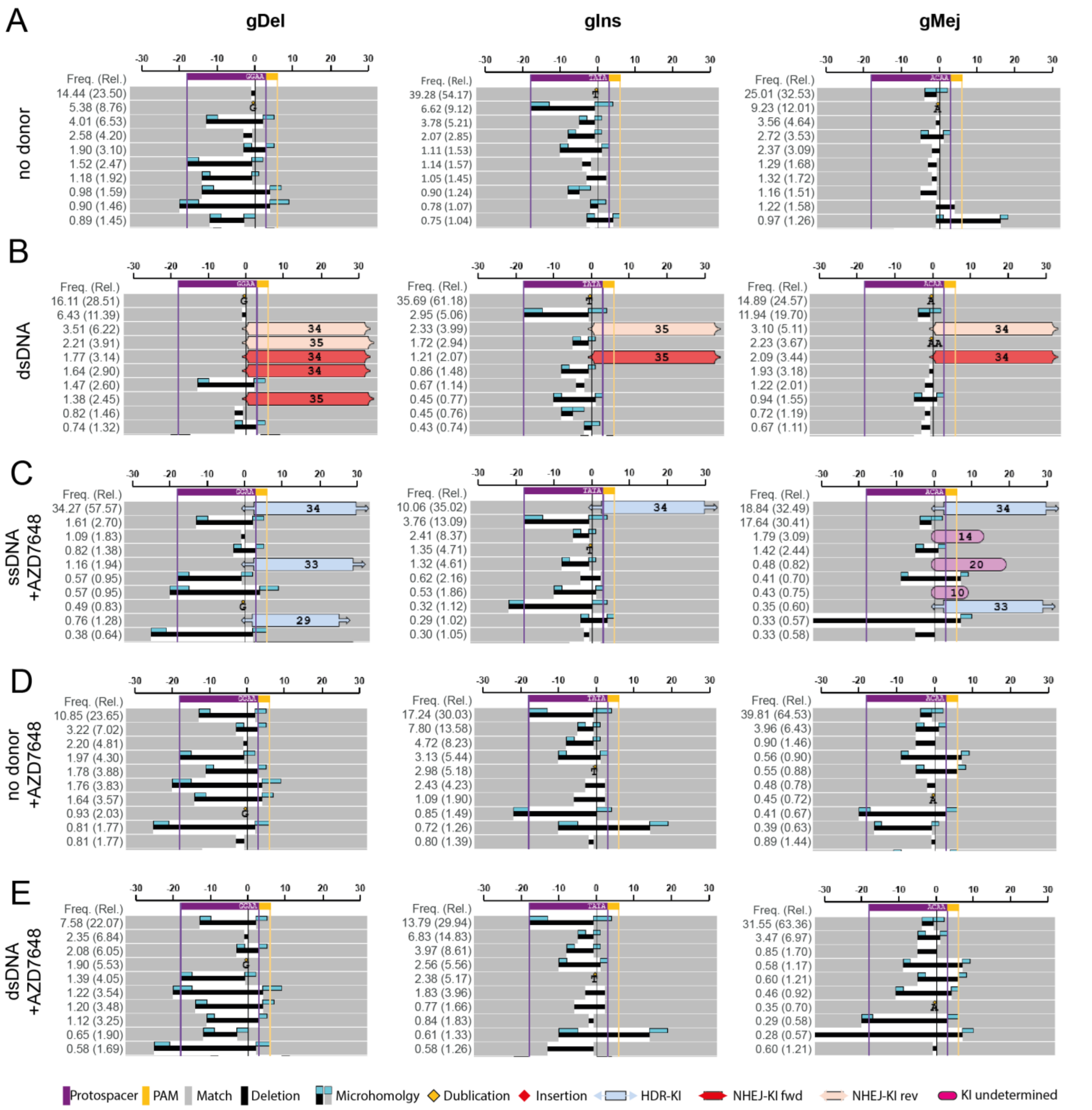

**Supplementary Figure S2 related to Figure 2: Graphical presentation of KI-Seq analysis to assess DNA repair outcomes at *Sp*Cas9 induced DSBs.** Representative InDel profiles +/-30 bp around the cut site from *Sp*Cas9 edited HEK293T cells at indicated loci, (**A-D**) without DNA donor, (**B,D**) with dsDNA (**C**) or ssDNA donor. Cells were either treated with (**A,B**) DMSO or (**C-E**) 1 µM AZD7648. Each variant is annotated with absolute frequencies (Freq.) = fraction of mapped reads and relative frequencies (Rel.) in brackets = fraction of mutated reads.

**Supplementary Figure S3.**

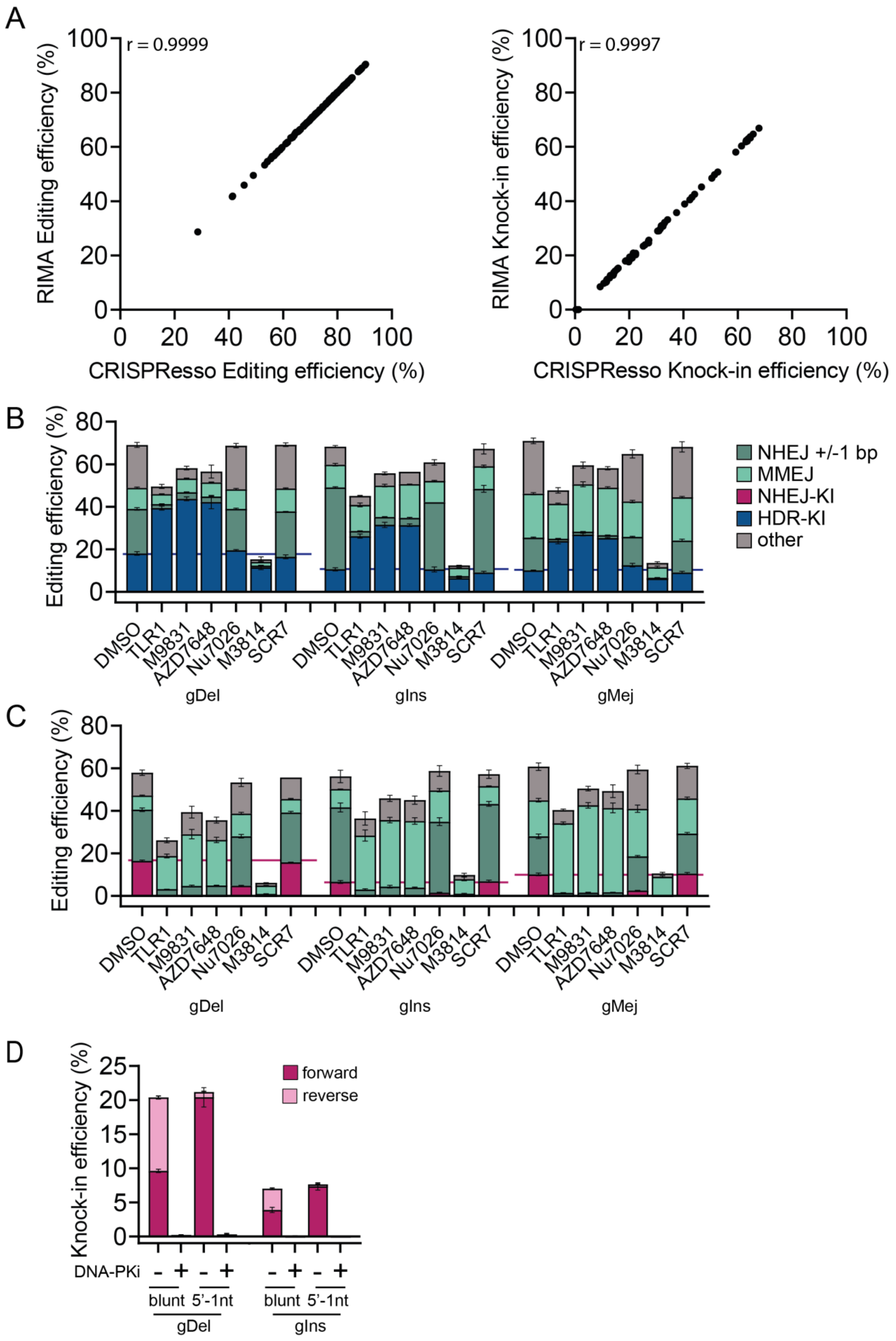

**Supplementary Figure S3 related to Figure 2: KI-Seq determines DNA repair outcomes at *Sp*Cas9 induced DSBs.** (**A**) Benchmarking KI-Seq to widely used CRISPResso2. Comparison of editing efficiencies and overall knock-in (perfect + imperfect knock-ins) of samples from several experiments (n=137). The dataset contains samples from three target sites, with and without ssDNA, and treatment with 1 µM AZD7648 or corresponding DMSO controls. Pearson correlation coefficient is shown. (**B,C**) Impact of NHEJ-repair pathway inhibitors on editing frequencies at three target sites (**B**) with the presence of ssDNA or (**C**) dsDNA analysed with KI-Seq. DNA-PK inhibitors from the TLR screen TLR1 (3 µM), M9831 (3 µM) and AZD7648 (3 µM) are compared to previously published compounds targeting DNA-PK Nu7026 (5.6 µM), M3814 (3 µM) or LIG4 SCR7 (10 µM). Bar graphs depict mean efficiencies ± SD (n=3, technical replicates) in mapped reads. (**D**) *Sp*Cas9 promotes directional NHEJ-mediated KI. Bar graphs show mean forward and reverse insertion efficiencies ± SD (n=3, technical replicates) for two types of dsDNA donor (blunt-ended and one nucleotide 5'-overhangs) at two target sites assessed with KI-Seq. Experiments were performed with 1 µM DNA-PK inhibitor (M9831) or DMSO control added 3 hours before plasmid transfections.

**Supplementary Figure S4.**

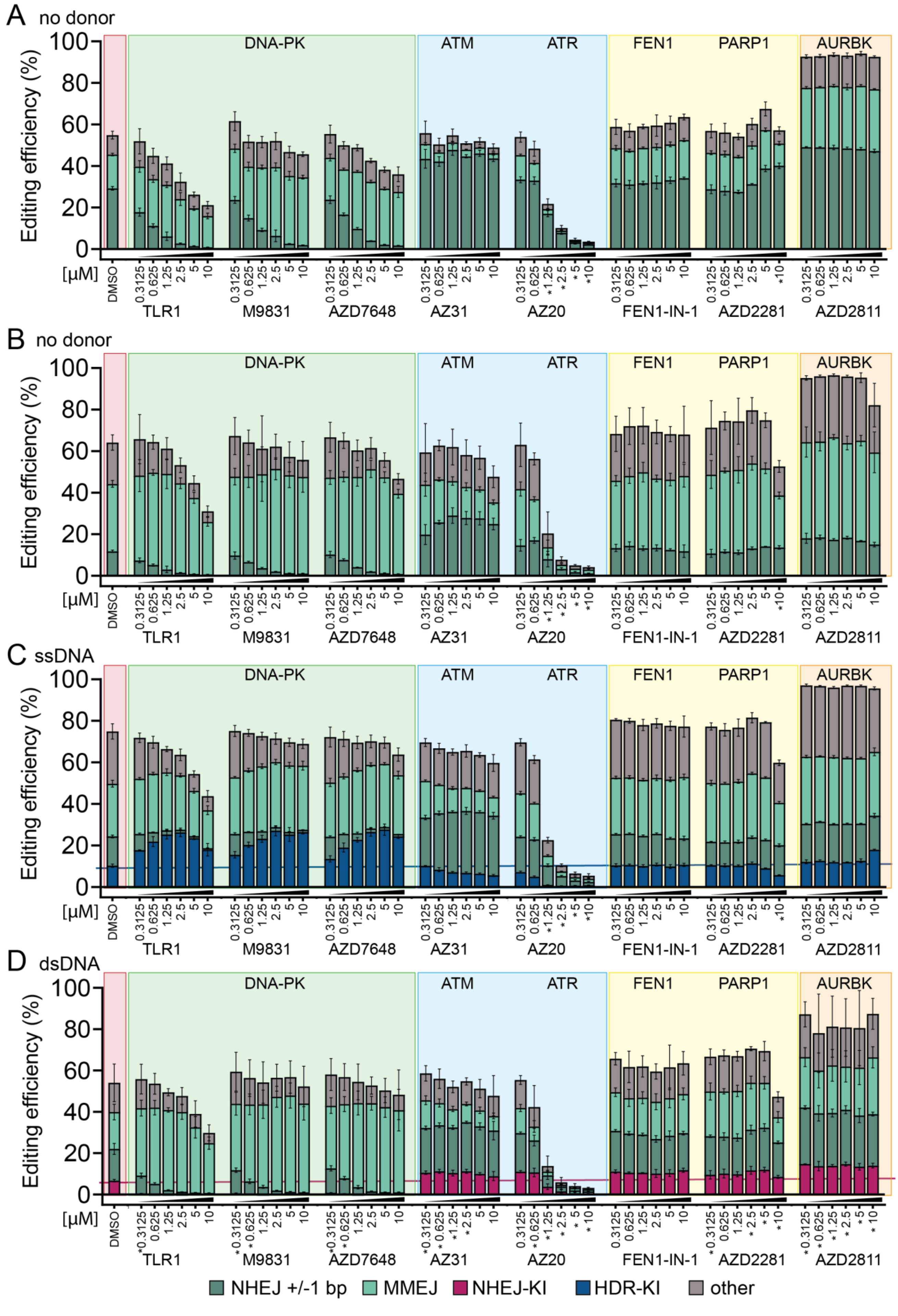

**Supplementary Figure S4 related to Figure 2: Extended analysis of the effect of TLR screen derived inhibitors on DNA repair outcomes at *Sp*Cas9 induced DSBs with KI-Seq** (**A-D**) Editing frequencies by repair pathway in HEK293T cells targeting (**A**) gIns or (**B-D**) gMej genomic loci, (**A,B**) without oligo donor, (**C**) with ssDNA or (**D**) dsDNA. KI-Seq was performed on cells treated with different concentrations of DNA repair inhibitors (0.3 – 10 µM) or DMSO control. Horizontal lines represent mean KI efficiencies in DMSO treated cells. Compounds and their associated targets are shown. Asterisk indicates treatments affecting cell confluency. Stacked bar graphs depict mean ± SD (n=3, technical replicates).

**Supplementary Figure S5.**

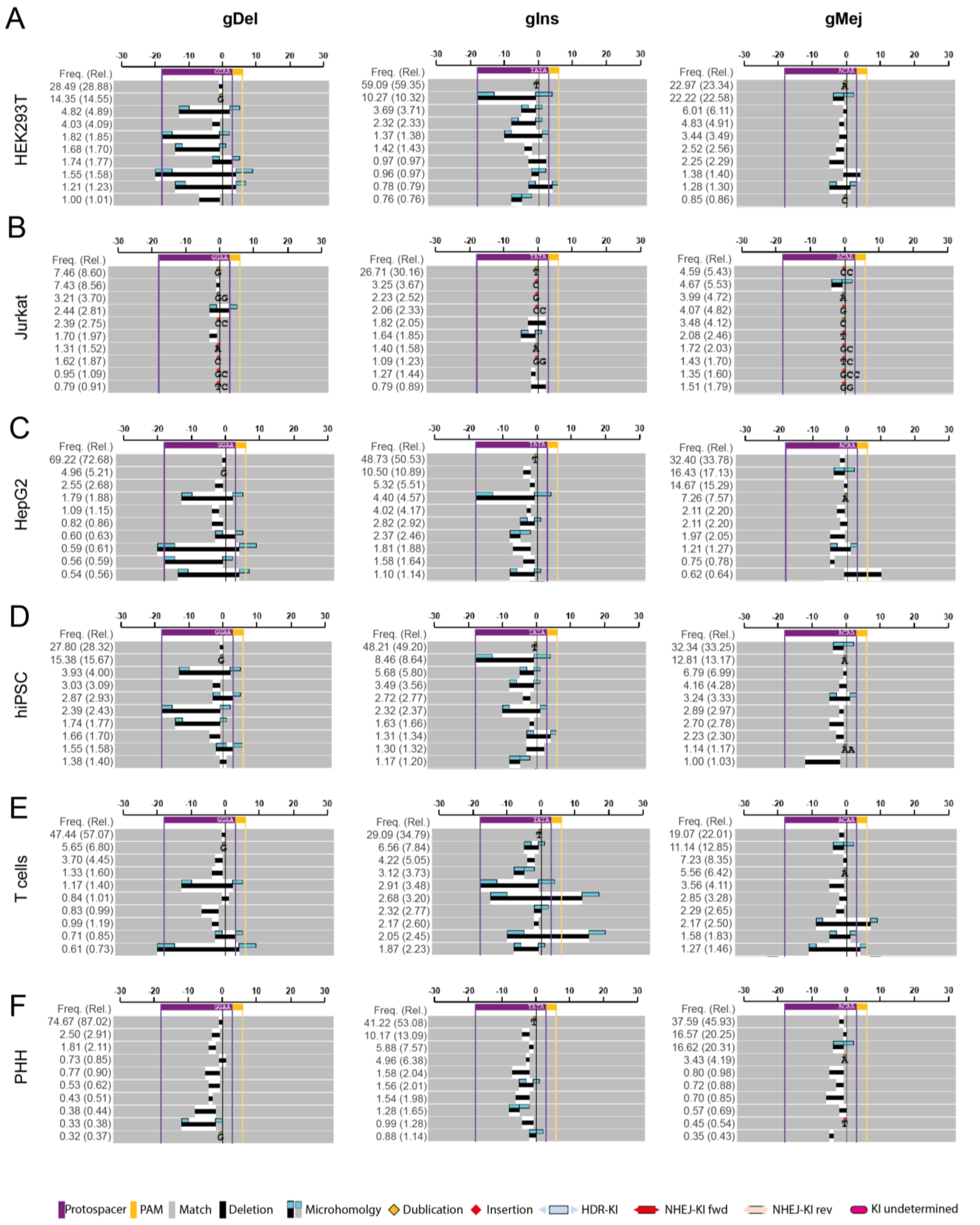

**Supplementary Figure S5 related to Figure 3: Mutation patterns can vary between different cell lines at the same target site.** (**A-F**) Representative InDel profiles +/-30 bp around the *Sp*Cas9 RNP-edited cut site at indicated loci and cell lines. Each variant is annotated with absolute frequencies (Freq.) = fraction of mapped reads and relative frequencies (Rel.) in brackets = fraction of mutated reads.

**Supplementary Figure S6.**

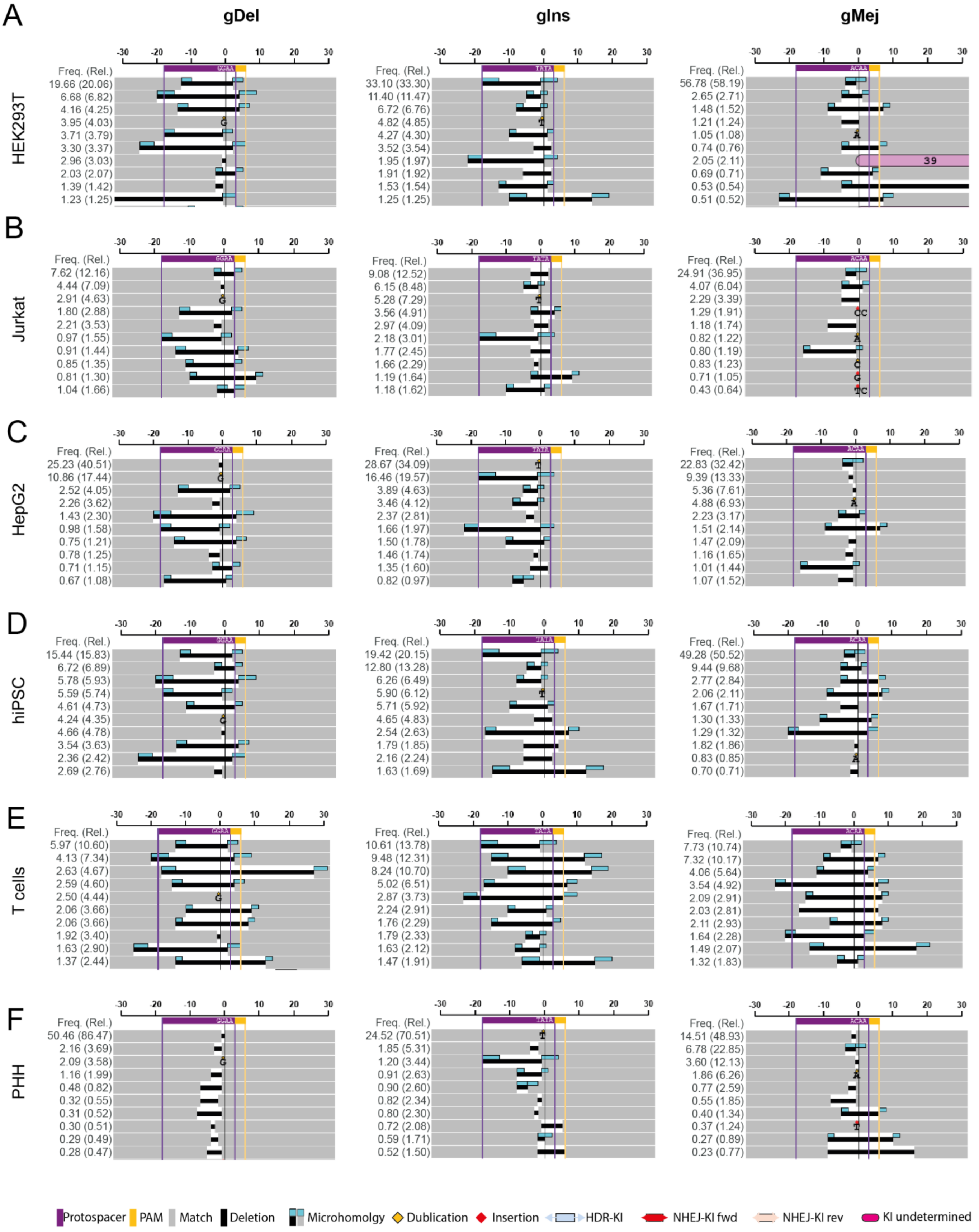

**Supplementary Figure S6 related to Figure 3: DNA-PK inhibition shifts mutation pattern towards longer deletions and increases the frequency of MMEJ-mediated deletions.** (**A-F**) Representative mutation patterns +/- 30 bp around the *Sp*Cas9 RNP-edited cut site at indicated loci and cell types in presence of 1 µM AZD7648. Each variant is annotated with absolute frequencies (Freq.) = fraction of mapped reads and relative frequencies (Rel.) in brackets = fraction of mutated reads.

**Supplementary Figure S7.**

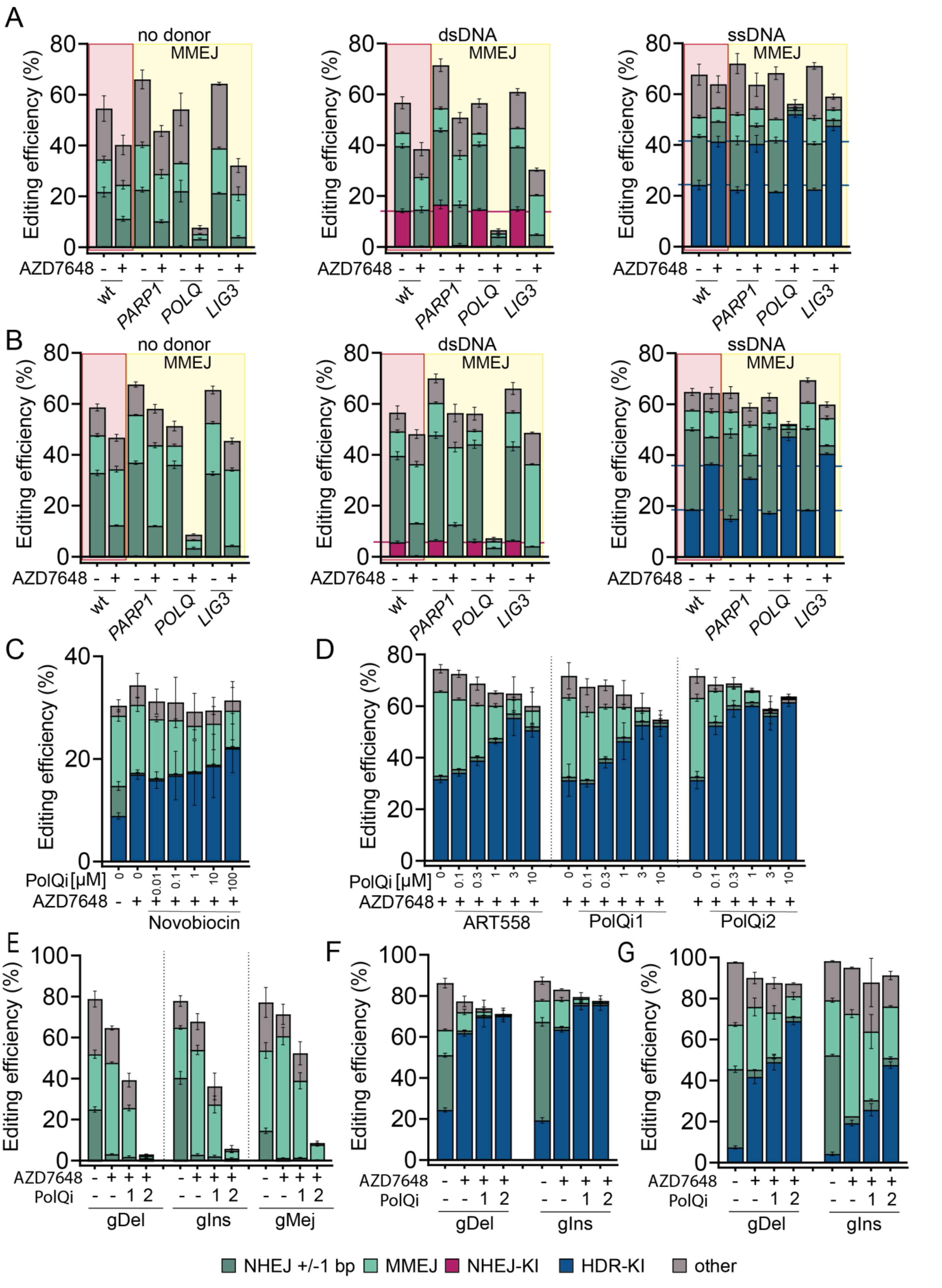

**Supplementary Figure S7 related to Figure 4: Simultaneous PolΘ and DNA-PK inhibition increases frequency and precision of ssDNA-mediated integration.** (**A,B**) Editing efficiencies of different repair events at (**A**) gDel and (**B**) gIns target sites with depicted DNA donors for HEK293T wild-type (wt) cells and three knockout cell pools of enzymes involved in MMEJ repair. Cells were treated with 1 µM AZD7648 or DMSO control and transfected with plasmids and ssDNA 3 hours later. Horizontal lines illustrate the mean knock-in efficiency in DMSO or AZD7648 treated wt cells. Bar graphs represent mean values ± SD (n=3, technical replicates). (**C**) Editing efficiencies of different repair events in HEK239T cells, co-treated with 1 µM AZD7648 and increasing concentrations of novobiocin (0.01 – 100 µM), or DMSO control. Cells were treated with compounds and transfected with plasmids and ssDNA 1-3 hours later. Bar graphs show mean values ± SD (n=3, technical replicates). (**D**) Dose-dependent effect on distribution of DNA repair events in gMej-targeted HEK293T cells treated with several PolΘ inhibitors (0.1 – 10 µM) in the background of 1 µM AZD7648 1-3 hours before plasmid and ssDNA transfection. Bar graphs illustrate mean editing efficiencies ± SD (n=3, technical replicates). (**E**) Editing efficiencies of different repair events at selected sites in HEK293T cells co-transfected with *Sp*Cas9 and sgRNA plasmids. Experiments were performed with 1-3 hours of pre-treatment using DMSO, 1 µM AZD7648, and 1 µM AZD7648 in combination with 3 µM PolQi1 or PolQi2. Bar graphs represent mean editing efficiencies ± SD (n=3, biological replicates). (**F**) Frequencies of different repair events at gDel and gIns target sites in HEK293T cells co-transfected with ssDNA, *Sp*Cas9 and sgRNA plasmids. Cells were treated 1-3 hours before transfections with DMSO, 1 µM DNA-PK inhibitor AZD7648, and 1 µM AZD7648 in combination with 3 µM PolΘ inhibitor 1 or 2. Bar graphs represent mean editing efficiencies ± SD (n=3, biological replicates). (**G**) Bar graphs depict mean editing efficiencies of different repair events ± SD (n=3, technical replicates) in *Sp*Cas9-inducible hiPSC transfected with sgRNA gDel and gIns in presence of DMSO, 1 µM AZD7648, and 1 µM AZD7648 in combination with 3 µM PolQi1 or PolQi2.

**Supplementary Figure S8.**

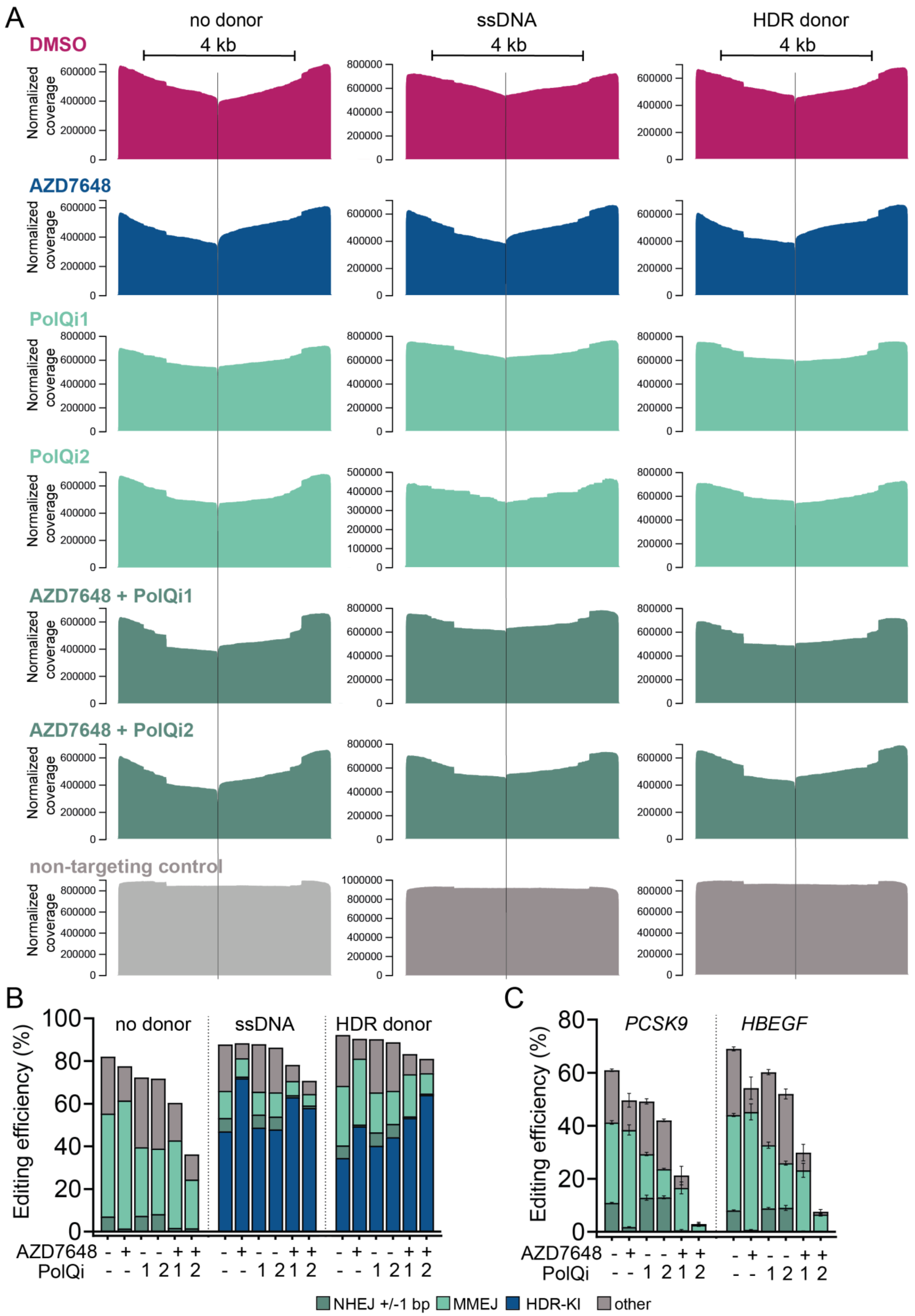

**Supplementary Figure S8 related to Figure 5: 2iHDR retains large deletions frequencies and decreases translocations.** (**A**) Coverage of long-range sequencing reads aligned to the edited *HBEGF* locus normalized to counts per million reads. Treatments with DMSO or small molecules (1 µM AZD7648, 3 µM PolQi1 and 3 µM PolQi2) and knock-in donors were used as depicted. (**B**) Editing efficiencies of different repair events for samples used in large deletion experiments targeting *HBEGF* with DNA donors depicted, analysed with KI-Seq. Cells were treated with indicated inhibitors (1 µM AZD7648, 3 µM PolQi1 and 3 µM PolQi2) or DMSO control (n=1). (**C**) Bar graphs show mean editing efficiencies of different repair events for samples used in translocation experiments (n=3, technical replicates). KI-Seq was performed for the indicated loci in presence of DMSO control or small molecules (1 µM AZD7648, 3 µM PolQi1, or 3 µM PolQi2).

**Supplementary Figure S9.**

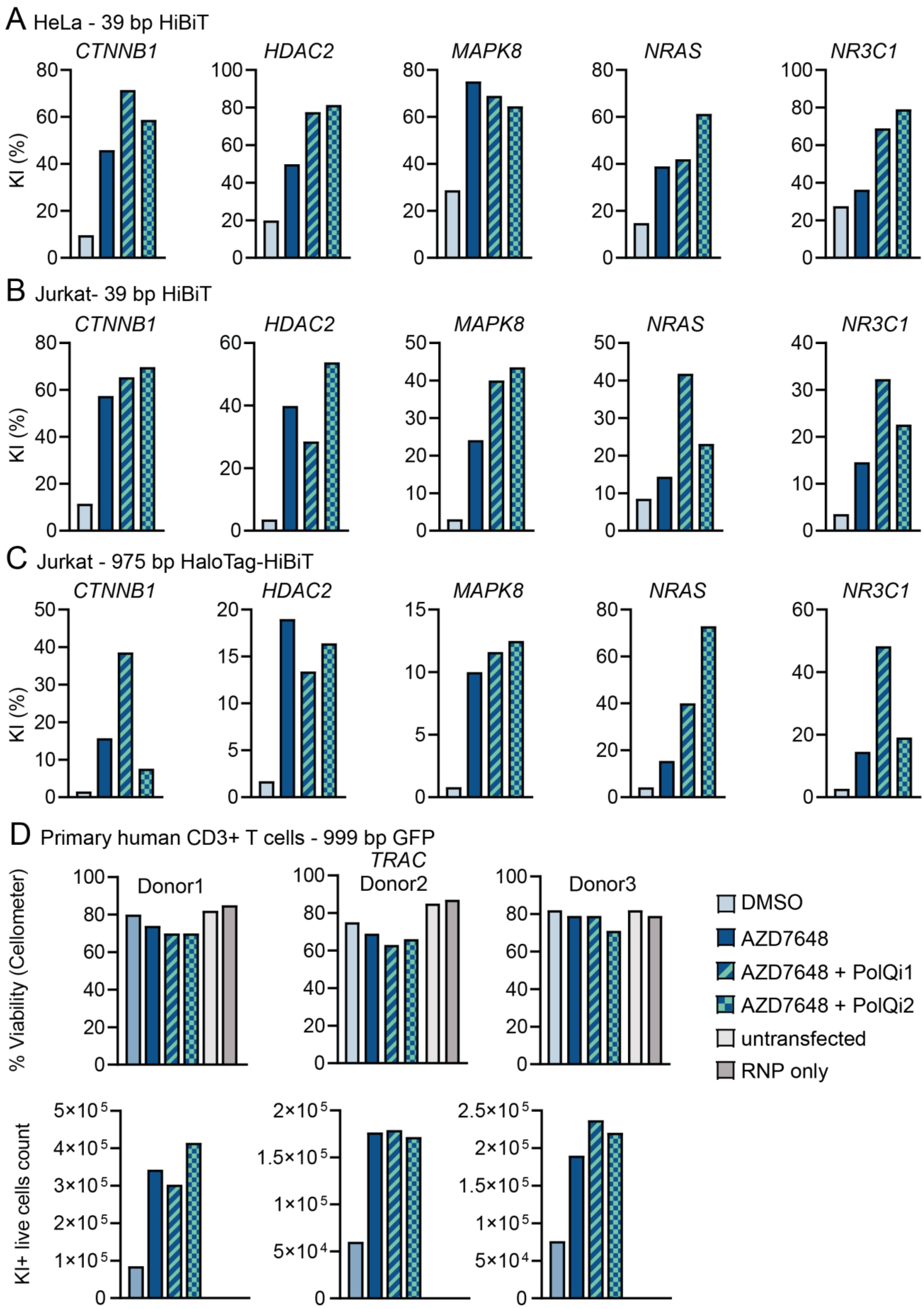

**Supplementary Figure S9 related to Figure 6: 2iHDR increases integration of different DNA donor templates with various integration-to-cut-site distances in diverse cell types. Validation of integration with droplet digital PCR (ddPCR) and cell viability measurement.** (**A-C**) Quantification of HiBiT and HaloTag-HiBiT integration assessed with ddPCR at indicated genomic loci and cell types. Treatment with 1 µM AZD7648 is compared to 2iHDR treatment (1 µM AZD7648 and 3 µM PolQi1 or PolQi2) or DMSO control. Bar graphs represent integration efficiency in percent (n=1). (**D**) Bar graphs show percentage viable CD3+ T cells assessed with trypan blue (upper panel) and corresponding live cell counts of knock-in containing cells (KI+) (lower panel). Cells were treated with DMSO, 1 µM AZD7648 or a combination of 1 µM AZD7648 and 3 µM PolQi1 or PolQi2 and compared to untreated or RNP controls as indicated.

**Supplementary Figure S10.**

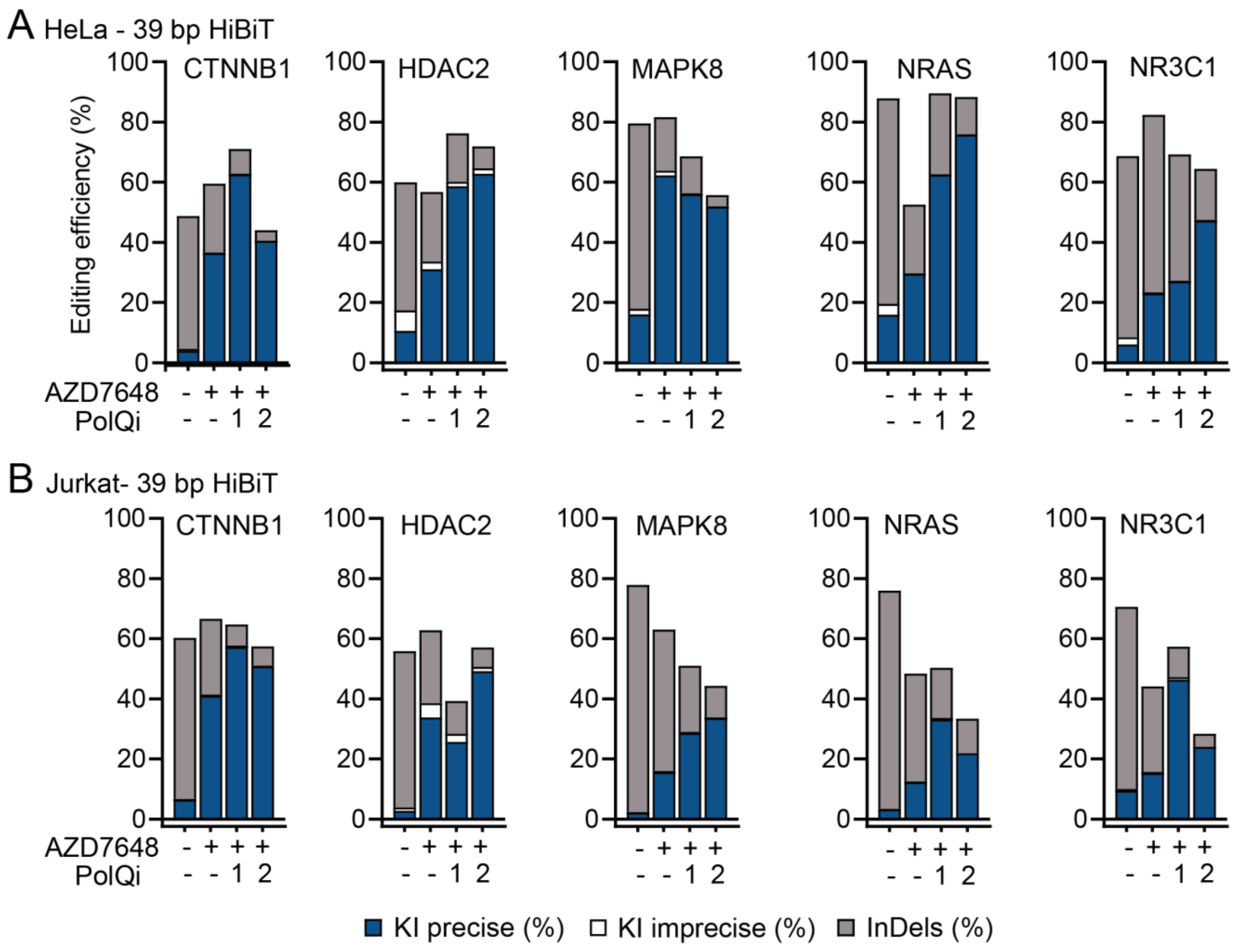

**Supplementary Figure S10 related to Figure 6: 2iHDR increases integration of different DNA donor templates with various integration-to-cut-site distances in diverse cell types. Deep-targeted amplicon sequencing.** (**A,B**) Representative editing efficiencies of different repair events in indicated cell types at various target sites evaluated with CRISPResso2 quantification window 1 (n=1). Knock-in precise = perfect HiBiT insertion, knock-in imprecise = HiBiT integration with additional mutations, InDels = all additional mutations independent from HiBiT integration. Experiments compare treatments with DMSO control, 1 µM AZD7648 and 2iHDR (1 µM AZD7648 and 3 µM PolQi1 or PolQi2).

**Supplementary Data S1.** Plasmids used in this study.

CMV-SpCas9-2AGFP

CCCGTAGAAAAGATCAAAGGATCTTCTTGAGATCCTTTTTTTCTGCGCGTAATCTGCTGCTTGCAAACAAAAAAACCACCGCTACCAGCGGTG
GTTTGTTTGCCGGATCAAGAGCTACCAACTCTTTTTCCGAAGGTAACTGGCTTCAGCAGAGCGCAGATACCAAATACTGTCCTTCTAGTGTAG
CCGTAGTTAGGCCACCACTTCAAGAACTCTGTAGCACCGCCTACATACCTCGCTCTGCTAATCCTGTTACCAGTGGCTGCTGCCAGTGGCGAT
AAGTCGTGTCTTACCGGGTTGGACTCAAGACGATAGTTACCGGATAAGGCGCAGCGGTCGGGCTGAACGGGGGGTTCGTGCACACAGCCCAGC
TTGGAGCGAACGACCTACACCGAACTGAGATACCTACAGCGTGAGCTATGAGAAAGCGCCACGCTTCCCGAAGGGAGAAAGGCGGACAGGTAT
CCGGTAAGCGGCAGGGTCGGAACAGGAGAGCGCACGAGGGAGCTTCCAGGGGGAAACGCCTGGTATCTTTATAGTCCTGTCGGGTTTCGCCAC
CTCTGACTTGAGCGTCGATTTTTGTGATGCTCGTCAGGGGGGCGGAGCCTATGGAAAAACGCCAGCAACGCGGCCTTTTTACGGTTCCTGGGC
TTTTGCTGGCCTTTTGCTCACATGTTCTTGCTGCTTCGCGATGTACGGGCCAGATATACGCGTTGACATTGATTATTGACTAGTTATTAATAG
TAATCAATTACGGGGTCATTAGTTCATAGCCCATATATGGAGTTCCGCGTTACATAACTTACGGTAAATGGCCCGCCTGGCTGACCGCCCAAC
GACCCCCGCCCATTGACGTCAATAATGACGTATGTTCCCATAGTAACGCCAATAGGGACTTTCCATTGACGTCAATGGGTGGACTATTTACGG
TAAACTGCCCACTTGGCAGTACATCAAGTGTATCATATGCCAAGTACGCCCCCTATTGACGTCAATGACGGTAAATGGCCCGCCTGGCATTAT
GCCCAGTACATGACCTTATGGGACTTTCCTACTTGGCAGTACATCTACGTATTAGTCATCGCTATTACCATGGTGATGCGGTTTTGGCAGTAC
ATCAATGGGCGTGGATAGCGGTTTGACTCACGGGGATTTCCAAGTCTCCACCCCATTGACGTCAATGGGAGTTTGTTTTGGCACCAAAATCAA
CGGGACTTTCCAAAATGTCGTAACAACTCCGCCCCATTGACGCAAATGGGCGGTAGGCGTGTACGGTGGGAGGTCTATATAAGCAGAGCTCTC
TGGCTAACTAGAGAACCCACTGCTTACTGGCTTATCGAAATTAATACGACTCACTATAGGGAGACCCAAGCTGGCTAGCGTTTAAACTTAAGC
TGATCCACTAGTCCAGTGTGGTGGAATTCGCCATGGCTCCTAAGAAAAAGCGGAAGGTGGACAAGAAATACTCAATCGGGCTGGACATCGGAA
CTAACTCAGTGGGGTGGGCAGTCATTACTGACGAGTACAAAGTGCCAAGCAAGAAATTTAAGGTCCTGGGCAACACCGATAGGCACTCCATCA
AGAAAAATCTGATTGGGGCCCTGCTGTTCGACTCTGGAGAGACAGCTGAAGCAACTAGACTGAAAAGGACTGCTAGAAGGCGCTATACCCGGC
GAAAGAATCGCATCTGCTACCTGCAGGAGATTTTCTCTAACGAAATGGCCAAGGTGGACGATAGTTTCTTTCATCGGCTGGAGGAATCATTCC
TGGTCGAGGAAGATAAGAAACACGAGAGACATCCTATCTTTGGAAACATTGTGGACGAGGTCGCTTATCACGAAAAATACCCCACCATCTATC
ATCTGCGCAAGAAACTGGTGGACTCTACAGATAAAGCAGACCTGCGGCTGATCTATCTGGCCCTGGCTCACATGATTAAGTTCAGAGGCCATT
TTCTGATCGAGGGAGATCTGAACCCAGACAATAGCGATGTGGACAAGCTGTTCATCCAGCTGGTCCAGACATACAATCAGCTGTTTGAGGAAA
ACCCTATTAATGCATCTGGCGTGGACGCAAAAGCCATCCTGAGTGCCAGGCTGTCTAAGAGTAGAAGGCTGGAGAACCTGATCGCTCAGCTGC
CAGGCGAAAAGAAAAACGGCCTGTTTGGAAATCTGATTGCACTGTCACTGGGACTGACACCTAACTTCAAGAGCAATTTTGATCTGGCCGAGG
ACGCTAAACTGCAGCTGAGCAAGGACACTTATGACGATGACCTGGATAACCTGCTGGCTCAGATCGGAGATCAGTACGCAGACCTGTTCCTGG
CCGCTAAGAATCTGTCTGACGCTATCCTGCTGAGTGATATTCTGCGGGTGAACACCGAGATTACAAAAGCCCCTCTGTCAGCTAGCATGATCA
AGAGATATGACGAGCACCATCAGGATCTGACCCTGCTGAAGGCACTGGTGCGCCAGCAGCTGCCCGAGAAGTACAAGGAAATCTTCTTTGATC
AGAGTAAGAACGGGTACGCCGGTTATATTGACGGCGGAGCTTCACAGGAGGAATTCTACAAGTTTATCAAACCTATTCTGGAGAAGATGGACG
GCACCGAGGAACTGCTGGTGAAACTGAATCGCGAGGACCTGCTGCGCAAGCAGCGGACATTTGATAACGGCTCCATCCCCCACCAGATTCATC
TGGGAGAGCTGCACGCAATCCTGCGACGACAGGAAGACTTCTACCCATTTCTGAAGGATAACCGCGAGAAGATCGAAAAAATTCTGACCTTCC
GGATCCCTTACTATGTGGGGCCCCTGGCAAGGGGTAATTCCCGCTTTGCCTGGATGACACGGAAATCTGAGGAAACAATCACTCCTTGGAACT
TCGAGGAAGTGGTCGATAAGGGAGCTTCCGCACAGTCTTTCATCGAGAGAATGACAAACTTCGACAAAAACCTGCCAAATGAGAAAGTGCTGC
CTAAGCACAGTCTGCTGTACGAGTATTTCACAGTCTATAACGAACTGACTAAGGTGAAATACGTCACCGAGGGGATGAGGAAGCCCGCCTTCC
TGAGCGGTGAACAGAAGAAAGCTATCGTGGACCTGCTGTTTAAAACCAATCGCAAGGTGACAGTCAAGCAGCTGAAGGAGGACTACTTCAAGA
AAATTGAATGTTTCGATTCTGTGGAGATCAGTGGCGTCGAAGACAGATTTAACGCTTCTCTGGGAACCTACCACGATCTGCTGAAGATCATTA
AGGATAAAGACTTCCTGGACAACGAGGAAAATGAGGATATCCTGGAAGACATTGTGCTGACCCTGACACTGTTTGAGGATCGCGAAATGATCG
AGGAACGGCTGAAAACTTATGCCCATCTGTTCGATGACAAGGTGATGAAACAGCTGAAGCGAAGAAGGTACACCGGCTGGGGACGACTGAGCA
GAAAGCTGATCAACGGCATTCGGGACAAACAGAGTGGAAAGACTATCCTGGACTTTCTGAAATCAGATGGCTTCGCTAACAGAAATTTTATGC
AGCTGATTCACGATGACAGCCTGACCTTCAAAGAGGATATCCAGAAGGCACAGGTGTCCGGGCAGGGTGACTCTCTGCACGAGCATATCGCAA
ACCTGGCCGGGTCCCCCGCCATCAAGAAAGGTATTCTGCAGACCGTGAAGGTGGTCGATGAGCTGGTGAAAGTCATGGGCAGGCATAAGCCAG
AAAACATCGTGATTGAGATGGCCCGCGAAAATCAGACCACACAGAAAGGACAGAAGAACAGCCGCGAGCGGATGAAAAGGATCGAGGAAGGCA
TTAAGGAACTGGGATCCCAGATCCTGAAAGAGCACCCTGTGGAAAACACTCAGCTGCAGAATGAGAAGCTGTATCTGTACTATCTGCAGAATG
GGCGGGATATGTACGTGGACCAGGAGCTGGATATTAACCGACTGTCTGATTACGACGTGGATCATATCGTCCCACAGTCATTCCTGAAAGATG
ACAGCATTGACAATAAGGTGCTGACCCGGAGTGACAAAAACCGAGGAAAGAGTGATAATGTCCCTTCAGAGGAAGTGGTCAAGAAAATGAAGA
ACTACTGGAGACAGCTGCTGAATGCCAAACTGATCACACAGCGAAAGTTTGATAACCTGACTAAAGCTGAGAGAGGGGGTCTGTCAGAACTGG
ACAAAGCAGGCTTCATCAAGCGACAGCTGGTGGAGACCAGACAGATCACAAAGCACGTCGCTCAGATTCTGGATAGCAGGATGAACACAAAGT
ACGATGAGAATGACAAACTGATCCGCGAAGTGAAGGTCATTACTCTGAAGTCAAAACTTGTGAGCGACTTCAGAAAGGATTTCCAGTTCTACA
AAGTCAGGGAGATCAACAATTATCACCATGCTCATGACGCATACCTGAACGCAGTGGTCGGGACCGCCCTGATTAAGAAATACCCCAAACTGG
AGAGCGAATTCGTGTACGGTGACTATAAGGTGTACGATGTCAGAAAAATGATCGCCAAGAGTGAGCAGGAAATTGGAAAAGCCACCGCTAAGT
ATTTCTTTTACTCAAACATCATGAATTTCTTTAAGACTGAGATCACCCTGGCAAATGGGGAAATCCGAAAGAGACCACTGATTGAGACTAACG
GCGAGACCGGAGAAATCGTGTGGGACAAGGGTAGGGATTTTGCCACAGTGCGCAAGGTCCTGTCCATGCCTCAAGTGAATATTGTCAAGAAAA
CAGAGGTGCAGACTGGCGGATTCAGTAAGGAATCAATTCTGCCCAAACGGAACTCTGATAAGCTGATCGCCCGAAAGAAAGACTGGGATCCCA
AGAAATATGGGGGTTTCGACTCCCCAACAGTGGCTTACTCTGTCCTGGTGGTCGCAAAGGTGGAGAAGGGGAAAAGCAAGAAACTGAAATCCG
TCAAGGAGCTGCTGGGTATCACTATTATGGAGAGGAGCTCCTTCGAGAAGAACCCCATCGATTTTCTGGAGGCTAAAGGCTATAAGGAAGTGA
AGAAAGACCTGATCATTAAACTGCCAAAGTACAGCCTGTTTGAGCTGGAAAACGGAAGGAAGCGAATGCTGGCATCCGCAGGAGAGCTGCAGA
AGGGTAATGAACTGGCCCTGCCTTCTAAGTACGTGAACTTCCTGTATCTGGCTAGCCACTACGAGAAGCTGAAAGGCTCCCCCGAGGATAACG
AACAGAAACAGCTGTTTGTGGAGCAGCACAAGCATTATCTGGACGAGATCATTGAACAGATTAGCGAGTTCTCCAAAAGAGTGATCCTGGCTG
ACGCAAATCTGGATAAGGTCCTGAGCGCATACAACAAACACAGAGATAAGCCAATCAGGGAGCAGGCCGAAAATATCATTCATCTGTTCACTC
TGACCAACCTGGGAGCCCCTGCAGCCTTCAAGTATTTTGACACTACCATCGATCGGAAACGATACACATCCACTAAGGAGGTGCTGGACGCTA
CCCTGATTCACCAGAGCATTACCGGCCTGTATGAAACAAGGATTGACCTGTCTCAGCTGGGGGGCGACCTCGAGGATGGGGACGAGGGCAGAG
GAAGTCTGCTAACATGCGGTGACGTCGAGGAGAATCCTGGCCCAGCACCGGGATCCATGGTGAGCAAGGGCGAGGAGCTGTTCACCGGGGTGG
TGCCCATCCTGGTCGAGCTGGACGGCGACGTAAACGGCCACAAGTTCAGCGTGTCCGGCGAGGGCGAGGGCGATGCCACCTACGGCAAGCTGA
CCCTGAAGTTCATCTGCACCACCGGCAAGCTGCCCGTGCCCTGGCCCACCCTCGTGACCACCTTCACCTACGGCGTGCAGTGCTTCGCCCGCT
ACCCCGACCACATGAAGCAGCACGACTTCTTCAAGTCCGCCATGCCCGAAGGCTACGTCCAGGAGCGCACCATCTTCTTCAAGGACGACGGCA
ACTACAAGACCCGCGCCGAGGTGAAGTTCGAGGGCGACACCCTGGTGAACCGCATCGAGCTGAAGGGCATCGACTTCAAGGAGGACGGCAACA
TCCTGGGGCACAAGCTGGAGTACAACTACAACAGCCACAAGGTCTATATCACCGCCGACAAGCAGAAGAACGGCATCAAGGTGAACTTCAAGA

CCCGCCACAACATCGAGGACGGCAGCGTGCAGCTCGCCGACCACTACCAGCAGAACACCCCCATCGGCGACGGCCCCGTGCTGCTGCCCGACA
ACCACTACCTGAGCACCCAGTCCGCCCTGAGCAAAGACCCCAACGAGAAGCGCGATCACATGGTCCTGCTGGAGTTCGTGACCGCCGCCGGGA
TCACTCTCGGCATGGACGAGCTGTACAAGTGATAATCTAGAGGGCCCGTTTAAACCCGCTGATCAGCCTCGACTGTGCCTTCTAGTTGCCAGC
CATCTGTTGTTTGCCCCTCCCCCGTGCCTTCCTTGACCCTGGAAGGTGCCACTCCCACTGTCCTTTCCTAATAAAATGAGGAAATTGCATCGC
ATTGTCTGAGTAGGTGTCATTCTATTCTGGGGGGTGGGGTGGGGCAGGACAGCAAGGGGGAGGATTGGGAAGACAATAGCAGGCATGCTGGGG
ATGCGGTGGGCTCTATGGCTTCTACTGGGCGGTTTTATGGACAGCAAGCGAACCGGAATTGCCAGCTGGGGCGCCCTCTGGTAAGGTTGGGAA
GCCCTGCAAAGTAAACTGGATGGCTTTCTCGCCGCCAAGGATCTGATGGCGCAGGGGATCAAGCTCTGATCAAGAGACAGGATGAGGATCGTT
TCGCATGATTGAACAAGATGGATTGCACGCAGGTTCTCCGGCCGCTTGGGTGGAGAGGCTATTCGGCTATGACTGGGCACAACAGACAATCGG
CTGCTCTGATGCCGCCGTGTTCCGGCTGTCAGCGCAGGGGCGCCCGGTTCTTTTTGTCAAGACCGACCTGTCCGGTGCCCTGAATGAACTGCA
AGACGAGGCAGCGCGGCTATCGTGGCTGGCCACGACGGGCGTTCCTTGCGCAGCTGTGCTCGACGTTGTCACTGAAGCGGGAAGGGACTGGCT
GCTATTGGGCGAAGTGCCGGGGCAGGATCTCCTGTCATCTCACCTTGCTCCTGCCGAGAAAGTATCCATCATGGCTGATGCAATGCGGCGGCT
GCATACGCTTGATCCGGCTACCTGCCCATTCGACCACCAAGCGAAACATCGCATCGAGCGAGCACGTACTCGGATGGAAGCCGGTCTTGTCGA
TCAGGATGATCTGGACGAAGAGCATCAGGGGCTCGCGCCAGCCGAACTGTTCGCCAGGCTCAAGGCGAGCATGCCCGACGGCGAGGATCTCGT
CGTGACCCATGGCGATGCCTGCTTGCCGAATATCATGGTGGAAAATGGCCGCTTTTCTGGATTCATCGACTGTGGCCGGCTGGGTGTGGCGGA
CCGCTATCAGGACATAGCGTTGGCTACCCGTGATATTGCTGAAGAGCTTGGCGGCGAATGGGCTGACCGCTTCCTCGTGCTTTACGGTATCGC
CGCTCCCGATTCGCAGCGCATCGCCTTCTATCGCCTTCTTGACGAGTTCTTCTGAATTATTAACGCTTACAATTTCCTGATGCGGTATTTTCT
CCTTACGCATCTGTGCGGTATTTCACACCGCATACAGGTGGCACTTTTCGGGGAAATGTGCGCGGAACCCCTATTTGTTTATTTTTCTAAATA
CATTCAAATATGTATCCGCTCATGAGACAATAACCCTGATAAATGCTTCAATAATAGCACGTGCTAAAACTTCATTTTTAATTTAAAAGGATC
TAGGTGAAGATCCTTTTTGATAATCTCATGACCAAAATCCCTTAACGTGAGTTTTCGTTCCACTGAGCGTCAGAC

### pMLU sgRNA backbone

TGTACAAAAAAGCAGGCTTTAAAGGAACCAATTCAGTCGACTGGATCCGGTACCAAGGTCGGGCAGGAAGAGGGCCTATTTCCCATGATTCCT
TCATATTTGCATATACGATACAAGGCTGTTAGAGAGATAATTAGAATTAATTTGACTGTAAACACAAAGATATTAGTACAAAATACGTGACGT
AGAAAGTAATAATTTCTTGGGTAGTTTGCAGTTTTAAAATTATGTTTTAAAATGGACTATCATATGCTTACCGTAACTTGAAAGTATTTCGAT
TTCTTGGCTTTATATATCTTGTGGAAAGGACGAAACACCGTATCGCAGGTGATTCTCTAGACATCATTAATTCCTAATTTTTGTTGACACTCT
ATCATTGATAGAGTTATTTTACCACTCCCTATCAGTGATAGAGAAAAGTGAAATGGCAACTGTTAACCAGCTGGTGCGCAAACCGCGTGCGCG
TAAGGTTGCAAAAAGCAACGTTCCTGCTCTGGAAGCTTGCCCGCAGAAGCGTGGTGTATGTACTCGCGTATATACCACCACCCCTAAAAAACC
AAACTCTGCACTGCGTAAAGTTTGTCGTGTGCGTCTGACTAACGGCTTCGAAGTTACCTCTTACATCGGTGGCGAAGGCCACAACCTGCAAGA
ACACTCCGTAATTCTGATCCGTGGTGGCCGTGTAAAAGACCTCCCAGGTGTGCGTTACCACACTGTTCGCGGTGCACTGGACTGTTCCGGTGT
TAAAGACCGTAAACAAGCGCGTTCTAAGTACGGTGTGAAGCGTCCAAAAGCTTAAAGGAGGACAATCATGGCCAAGCCTTTGTCTCAAGAAGA
ATCCACCCTCATTGAAAGAGCAACGGCTACAATCAACAGCATCCCCATCTCTGAAGACTACAGCGTCGCCAGCGCAGCTCTCTCTAGCGACGG
CCGCATCTTCACTGGTGTCAATGTATATCATTTTACTGGGGGACCTTGTGCAGAACTCGTGGTGCTGGGCACTGCTGCTGCTGCGGCAGCTGG
CAACCTGACTTGTATCGTCGCGATCGGAAATGAGAACAGGGGCATCTTGAGCCCCTGCGGACGGTGTCGACAGGTGCTTCTCGATCTGCATCC
TGGGATCAAAGCGATAGTGAAGGACAGTGATGGACAGCCGACGGCAGTTGGGATTCGTGAATTGCTGCCCTCTGGTTATGTGTGGGAGGGCTA
ACACCTGCATCCGTTTTAGAGCTAGAAATAGCAAGTTAAAATAAGGCTAGTCCGTTATCAACTTGAAAAAGTGGCACCGAGTCGGTGCTTTTT
TTCTAGACCCAGCTTTCTTGTACAAAGTTGGCATTAACGCGTTGACATTGATTATTGACTAGCCACTACGAGAAGCTGAAAGGCTCCCCCGAG
GATAACGAACAGAAACAGCTGTTTGTGGAGCAGCACAAGCATTATCTGGACGAGATCATTGAACAGATTAGCGAGTTCTCCAAAAGAGTGATC
CTGGCTGACGCAAATCTGGATAAGGTCCTGAGCGCATACAACAAACACAGAGATAAGCCAATCAGGGAGCAGGCCGAAAATATCATTCATCTG
TTCACTCTGACCAACCTGGGAGCCCCTGCAGCCTTCAAGTATTTTGACACTACCATCGATCGGAAACGATACACATCCACTAAGGAGGTGCTG
GACGCTACCCTGATTCACCAGAGCATTACCGGCCTGTATGAAACAAGGATTGACCTGTCTCAGCTGGGCGGCGACAGCAGGGCTGACCCCAAG
AAGAAGAGGAAGGTGTGATAACTCGAGTCTAGAGGGCCCGTTTAAACCCGCTGATCAGCCTCGACTGTGCCTTCTAGTTGCCAGCCATCTGTT
GTTTGCCCCTCCCCCGTGCCTTCCTTGACCCTGGAAGGTGCCACTCCCACTGTCCTTTCCTAATAAAATGAGGAAATTGCATCGCATTGTCTG
AGTAGGTGTCATTCTATTCTGGGGGGTGGGGTGGGGCAGGACAGCAAGGGGGAGGATTGGGAAGACAATAGCAGGCATGCTGGGGATGCGGTG
GGCTCTATGGCTTCTACTGGGCGGTTTTATGGACAGCAAGCGAACCGGAATTGCCAGCTGGGGCGCCCTCTGGTAAGGTTGGGAAGCCCTGCA
AAGTAAACTGGATGGCTTTCTCGCCGCCAAGGATCTGATGGCGCAGGGGATCAAGCTCTGATCAAGAGACAGGATGAGGATCGTTTCGCATGA
TTGAACAAGATGGATTGCACGCAGGTTCTCCGGCCGCTTGGGTGGAGAGGCTATTCGGCTATGACTGGGCACAACAGACAATCGGCTGCTCTG
ATGCCGCCGTGTTCCGGCTGTCAGCGCAGGGGCGCCCGGTTCTTTTTGTCAAGACCGACCTGTCCGGTGCCCTGAATGAACTGCAAGACGAGG
CAGCGCGGCTATCGTGGCTGGCCACGACGGGCGTTCCTTGCGCAGCTGTGCTCGACGTTGTCACTGAAGCGGGAAGGGACTGGCTGCTATTGG
GCGAAGTGCCGGGGCAGGATCTCCTGTCATCTCACCTTGCTCCTGCCGAGAAAGTATCCATCATGGCTGATGCAATGCGGCGGCTGCATACGC
TTGATCCGGCTACCTGCCCATTCGACCACCAAGCGAAACATCGCATCGAGCGAGCACGTACTCGGATGGAAGCCGGTCTTGTCGATCAGGATG
ATCTGGACGAAGAGCATCAGGGGCTCGCGCCAGCCGAACTGTTCGCCAGGCTCAAGGCGAGCATGCCCGACGGCGAGGATCTCGTCGTGACCC
ATGGCGATGCCTGCTTGCCGAATATCATGGTGGAAAATGGCCGCTTTTCTGGATTCATCGACTGTGGCCGGCTGGGTGTGGCGGACCGCTATC
AGGACATAGCGTTGGCTACCCGTGATATTGCTGAAGAGCTTGGCGGCGAATGGGCTGACCGCTTCCTCGTGCTTTACGGTATCGCCGCTCCCG
ATTCGCAGCGCATCGCCTTCTATCGCCTTCTTGACGAGTTCTTCTGAATTATTAACGCTTACAATTTCCTGATGCGGTATTTTCTCCTTACGC
ATCTGTGCGGTATTTCACACCGCATACAGGTGGCACTTTTCGGGGAAATGTGCGCGGAACCCCTATTTGTTTATTTTTCTAAATACATTCAAA
TATGTATCCGCTCATGAGACAATAACCCTGATAAATGCTTCAATAATAGCACGTGCTAAAACTTCATTTTTAATTTAAAAGGATCTAGGTGAA
GATCCTTTTTGATAATCTCATGACCAAAATCCCTTAACGTGAGTTTTCGTTCCACTGAGCGTCAGACCCCGTAGAAAAGATCAAAGGATCTTC
TTGAGATCCTTTTTTTCTGCGCGTAATCTGCTGCTTGCAAACAAAAAAACCACCGCTACCAGCGGTGGTTTGTTTGCCGGATCAAGAGCTACC
AACTCTTTTTCCGAAGGTAACTGGCTTCAGCAGAGCGCAGATACCAAATACTGTCCTTCTAGTGTAGCCGTAGTTAGGCCACCACTTCAAGAA
CTCTGTAGCACCGCCTACATACCTCGCTCTGCTAATCCTGTTACCAGTGGCTGCTGCCAGTGGCGATAAGTCGTGTCTTACCGGGTTGGACTC
AAGACGATAGTTACCGGATAAGGCGCAGCGGTCGGGCTGAACGGGGGGTTCGTGCACACAGCCCAGCTTGGAGCGAACGACCTACACCGAACT
GAGATACCTACAGCGTGAGCTATGAGAAAGCGCCACGCTTCCCGAAGGGAGAAAGGCGGACAGGTATCCGGTAAGCGGCAGGGTCGGAACAGG
AGAGCGCACGAGGGAGCTTCCAGGGGGAAACGCCTGGTATCTTTATAGTCCTGTCGGGTTTCGCCACCTCTGACTTGAGCGTCGATTTTTGTG
ATGCTCGTCAGGGGGGCGGAGCCTATGGAAAAACGCCAGCAACGCGGCCTTTTTACGGTTCCTGGGCTTTTGCTGGCCTTTTGCTCACATGTT
CTTGCTGCTTCGCGATGTACGGGCCAGATATACGCGT

### pMA sgRNA backbone

AGAGGGCCTATTTCCCATGATTCCTTCATATTTGCATATACGATACAAGGCTGTTAGAGAGATAATTAGAATTAATTTGACTGTAAACACAAA
GATATTAGTACAAAATACGTGACGTAGAAAGTAATAATTTCTTGGGTAGTTTGCAGTTTTAAAATTATGTTTTAAAATGGACTATCATATGCT
TACCGTAACTTGAAAGTATTTCGATTTCTTGGCTTTATATATCTTGTGGAAAGGACGAAACACCGNNNNNNNNNNNNNNNNNNNNGTTTTAGA
GCTAGAAATAGCAAGTTAAAATAAGGCTAGTCCGTTATCAACTTGAAAAAGTGGCACCGAGTCGGTGCTTTTTTTGCCTCATGGGCCTTCCGC
TCACTGCCCGCTTTCCAGTCGGGAAACCTGTCGTGCCAGCTGCATTAACATGGTCATAGCTGTTTCCTTGCGTATTGGGCGCTCTCCGCTTCC
TCGCTCACTGACTCGCTGCGCTCGGTCGTTCGGGTAAAGCCTGGGGTGCCTAATGAGCAAAAGGCCAGCAAAAGGCCAGGAACCGTAAAAAGG
CCGCGTTGCTGGCGTTTTTCCATAGGCTCCGCCCCCCTGACGAGCATCACAAAAATCGACGCTCAAGTCAGAGGTGGCGAAACCCGACAGGAC
TATAAAGATACCAGGCGTTTCCCCCTGGAAGCTCCCTCGTGCGCTCTCCTGTTCCGACCCTGCCGCTTACCGGATACCTGTCCGCCTTTCTCC
CTTCGGGAAGCGTGGCGCTTTCTCATAGCTCACGCTGTAGGTATCTCAGTTCGGTGTAGGTCGTTCGCTCCAAGCTGGGCTGTGTGCACGAAC
CCCCCGTTCAG

### TLR-Construct

ATCCTGTCCCTAGGCCACCCACTGTGGGGTCTGACCTCTTCTCTTCCTCCCACAGGGCCTCGAGAGATCTGGCAGCGGAGAGGGCAGAGGAAG
TCTTCTAACATGCGGTGACGTGGAGGAGAATCCCGGCCCTAGGCTCGAAATGACCGAGTACAAGCCCACGGTGCGCCTCGCCACCCGCGACGA
CGTCCCCCGGGCCGTACGCACCCTCGCCGCCGCGTTCGCCGACTACCCCGCCACGCGCCACACCGTCGACCCGGACCGCCACATCGAGCGGGT
CACCGAGCTGCAAGAACTCTTCCTCACGCGCGTCGGGCTCGACATCGGCAAGGTGTGGGTCGCGGACGACGGCGCCGCGGTGGCGGTCTGGAC
CACGCCGGAGAGCGTCGAAGCGGGGGCGGTGTTCGCCGAGATCGGCCCGCGCATGGCCGAGTTGAGCGGTTCCCGGCTGGCCGCGCAGCAACA
GATGGAAGGCCTCCTGGCGCCGCACCGGCCCAAGGAGCCCGCGTGGTTCCTGGCCACCGTCGGCGTCTCGCCCGACCACCAGGGCAAGGGTCT
GGGCAGCGCCGTCGTGCTCCCCGGAGTGGAGGCGGCCGAGCGCGCCGGGGTGCCCGCCTTCCTGGAGACCTCCGCGCCCCGCAACCTCCCCTT
CTACGAGCGGCTCGGCTTCACCGTCACCGCCGACGTCGAGGTGCCCGAAGGACCGCGCACCTGGTGCATGACCCGCAAGCCCGGTGCCCCTTC
CTAGACACATCCCTTATGGGAGATCTGGCAGCGGAGAGGGCAGAGGAAGTCTTCTAACATGCGGTGACGTGGAGGAGAATCCCGGCCCTAGGA
TGGCCAAGCCTTTGTCTCAAGAAGAATCCACCCTCATTGAAAGAGCAACGGCTACAATCAACAGCATCCCCATCTCTGAAGACTACAGCGTCG
CCAGCGCAGCTCTCTCTAGCGACGGCCGCATCTTCACTGGTGTCAATGTATATCATTTTACTGGGGGACCTTGTGCAGAACTCGTGGTGCTGG
GCACTGCTGCTGCTGCGGCAGCTGGCAACCTGACTTGTATCGTCGCGATCGGAAATGAGAACAGGGGCATCTTGAGCCCCTGCGGACGGTGCC
GACAGGTGCTTCTCGATCTGCATCCTGGGATCAAAGCCATAGTGAAGGACAGTGATGGACAGCCGACGGCAGTTGGGATTCGTGAATTGCTGC
CCTCTGGTTATGTGTGGGAGGGCTAACGCCCGCCCCACGACCCGCAGCGCCCGACCGAAAGGAGCGCACGACCCCATGGCTCCGACCGAAGCC
ACCCGGGGCGGCCCCGCCGACCCCGCACCCGCCCCCGAGGCCCACCGACTCTAGAGGGCCCGTTTAAACCCGCTGATCAGCCTCGACTGTGCC
TTCTAGTTGCCAGCCATCTGTTGTTTGCCCCTCCCCCGTGCCTTCCTTGACCCTGGAAGGTGCCACTCCCACTGTCCTTTCCTAATAAAATGA
GGAAATTGCATCGCATTGTCTGAGTAGGTGTCATTCTATTCTGGGGGGTGGGGTGGGGCAGGACAGCAAGGGGGAGGATTGGGAAGACAATAG
CAACCGGTGCTAGAATTCACTGGCCGTCGTTTTACAACGTCGTGACTGGGAAAACCCTGGCGTTACCCAACTTAATCGCCTTGCAGCACATCC
CCCTTTCGCCAGCTGGCGTAATAGCGAAGAGGCCCGCACCGATCGCCCTTCCCAACAGTTGCGCAGCCTGAATGGCGAATGGCGCCTGATGCG
GTATTTTCTCCTTACGCATCTGTGCGGTATTTCACACCGCATATGGTGCACTCTCAGTACAATCTGCTCTGATGCCGCATAGTTAAGCCAGCC
CCGACACCCGCCAACACCCGCTGACGCGCCCTGACGGGCTTGTCTGCTCCCGGCATCCGCTTACAGACAAGCTGTGACCGTCTCCGGGAGCTG
CATGTGTCAGAGGTTTTCACCGTCATCACCGAAACGCGCGAGACGAAAGGGCCTCGTGATACGCCTATTTTTATAGGTTAATGTCATGATAAT
AATGGTTTCTTAGACGTCAGGTGGCACTTTTCGGGGAAATGTGCGCGGAACCCCTATTTGTTTATTTTTCTAAATACATTCAAATATGTATCC
GCTCATGAGACAATAACCCTGATAAATGCTTCAATAATATTGAAAAAGGAAGAGTATGAGTATTCAACATTTCCGTGTCGCCCTTATTCCCTT
TTTTGCGGCATTTTGCCTTCCTGTTTTTGCTCACCCAGAAACGCTGGTGAAAGTAAAAGATGCTGAAGATCAGTTGGGTGCACGAGTGGGTTA
CATCGAACTGGATCTCAACAGCGGTAAGATCCTTGAGAGTTTTCGCCCCGAAGAACGTTTTCCAATGATGAGCACTTTTAAAGTTCTGCTATG
TGGCGCGGTATTATCCCGTATTGACGCCGGGCAAGAGCAACTCGGTCGCCGCATACACTATTCTCAGAATGACTTGGTTGAGTACTCACCAGT
CACAGAAAAGCATCTTACGGATGGCATGACAGTAAGAGAATTATGCAGTGCTGCCATAACCATGAGTGATAACACTGCGGCCAACTTACTTCT
GACAACGATCGGAGGACCGAAGGAGCTAACCGCTTTTTTGCACAACATGGGGGATCATGTAACTCGCCTTGATCGTTGGGAACCGGAGCTGAA
TGAAGCCATACCAAACGACGAGCGTGACACCACGATGCCTGTAGCAATGGCAACAACGTTGCGCAAACTATTAACTGGCGAACTACTTACTCT
AGCTTCCCGGCAACAATTAATAGACTGGATGGAGGCGGATAAAGTTGCAGGACCACTTCTGCGCTCGGCCCTTCCGGCTGGCTGGTTTATTGC
TGATAAATCTGGAGCCGGTGAGCGTGGGTCTCGCGGTATCATTGCAGCACTGGGGCCAGATGGTAAGCCCTCCCGTATCGTAGTTATCTACAC
GACGGGGAGTCAGGCAACTATGGATGAACGAAATAGACAGATCGCTGAGATAGGTGCCTCACTGATTAAGCATTGGTAACTGTCAGACCAAGT
TTACTCATATATACTTTAGATTGATTTAAAACTTCATTTTTAATTTAAAAGGATCTAGGTGAAGATCCTTTTTGATAATCTCATGACCAAAAT
CCCTTAACGTGAGTTTTCGTTCCACTGAGCGTCAGACCCCGTAGAAAAGATCAAAGGATCTTCTTGAGATCCTTTTTTTCTGCGCGTAATCTG
CTGCTTGCAAACAAAAAAACCACCGCTACCAGCGGTGGTTTGTTTGCCGGATCAAGAGCTACCAACTCTTTTTCCGAAGGTAACTGGCTTCAG
CAGAGCGCAGATACCAAATACTGTTCTTCTAGTGTAGCCGTAGTTAGGCCACCACTTCAAGAACTCTGTAGCACCGCCTACATACCTCGCTCT
GCTAATCCTGTTACCAGTGGCTGCTGCCAGTGGCGATAAGTCGTGTCTTACCGGGTTGGACTCAAGACGATAGTTACCGGATAAGGCGCAGCG
GTCGGGCTGAACGGGGGGTTCGTGCACACAGCCCAGCTTGGAGCGAACGACCTACACCGAACTGAGATACCTACAGCGTGAGCTATGAGAAAG
CGCCACGCTTCCCGAAGGGAGAAAGGCGGACAGGTATCCGGTAAGCGGCAGGGTCGGAACAGGAGAGCGCACGAGGGAGCTTCCAGGGGGAAA
CGCCTGGTATCTTTATAGTCCTGTCGGGTTTCGCCACCTCTGACTTGAGCGTCGATTTTTGTGATGCTCGTCAGGGGGGCGGAGCCTATGGAA
AAACGCCAGCAACGCGGCCTTTTTACGGTTCCTGGCCTTTTGCTGGCCTTTTGCTCACATGTTCTTTCCTGCGTTATCCCCTGATTCTGTGGA
TAACCGTATTACCGCCTTTGAGTGAGCTGATACCGCTCGCCGCAGCCGAACGACCGAGCGCAGCGAGTCAGTGAGCGAGGAAGCGGAAGAGCG
CCCAATACGCAAACCGCCTCTCCCCGCGCGTTGGCCGATTCATTAATGCAGCTGGCACGACAGGTTTCCCGACTGGAAAGCGGGCAGTGAGCG
CAACGCAATTAATGTGAGTTAGCTCACTCATTAGGCACCCCAGGCTTTACACTTTATGCTTCCGGCTCGTATGTTGTGTGGAATTGTGAGCGG
ATAACAATTTCACACAGGAAACAGCTATGACCATGATTACGCCAAGCTTCTTAAGTAGACGCGTCCATAGAGCCCACCGCATCCCCAGCATGC
CGATTTTACCACATTTGTAGAGGTTTTACTTGCTTTAAAAAACCTCCCACACCTCCCCCTGAACCTGAAACATAAAATGAATGCAATTGTTGT
TGTTAACTTGTTTATTGCAGCTTATAATGGTTACAAATAAAGCAATAGCATCACAAATTTCACAAATAAAGCATTTTTTTCACTGCATTCTAG
TTGTGGTTTGTCCAAACTCATCAATGTATCTTATCATGTCTGCTCGAAGCGGCCGCTACAGGAACAGGTGGTGGCGGCCCTCGGCGCGCTCGT
ACTGCTCCACGATGGTGTAGTCCTCGTTGTGGGAGGTGATGTCCAGCTTGGAGTCCACGTAGTAGTAGCCGGGCAGCTGCACGGGCTTCTTGG
CCATGTAGATGGACTTGAACTCCACCAGGTAGTGGCCGCCGTCCTTCAGCTTCAGGGCCTTGTGGATCTCGCCCTTCAGCACGCCGTCGCGGG
GGTACAGGCGCTCGGTGGAGGCCTCCCAGCCCATAGTCTTCTTCTGCATTACGGGGCCGTCGGAGGGGAAGTTCACGCCGATGAACTTCACCT
TGTAGATGAAGGAGCCGTCCTGCAGGGAGGAGTCCTGGGTCACGGTCACCACGCCGCCGTCCTCGAAGTTCATCACGCGCTCCCACTTGAAGC
CCTCGGGGAAGGACAGCTTCTTGTAGTCGGGGATGTCGGCGGGGTGCTTCACGTACACCTTGGAGCCGTACTGGAACTGGGGGGACAGGATGT
CCCAGGCGAAGGGCAGGGGGCCGCCCTTGGTCACCTTCAGCTTGGCGGTCTGGGTGCCCTCGTAGGGGCGGCCCTCGCCCTCGCCCTCGATCT
CGAACTCGTGGCCGTTCACGGAGCCCTCCATGCGCACCTTGAAGCGCATGAACTCCTTGATGACGTCCTCGGAGGAGGCAGGGCCCGGATTCT

CCTCCACGTCACCGCATGTTAGAAGACTTCCTCTGCCCTCACCGGGAGCTCTTACTTGTACAGCTCGTCCATGCCGAGAGTGATCCCGGCGGC
GGTCACGAACTCCAGCAGGACCATGTGATCGCGCTTCTCGTTGGGGTCTTTGCTCAGGGCGGACTGGGTGCTCAGGTAGTGGTTGTCGGGCAG
CAGCACGGGGCCGTCGCCAATAGGTGTGTTCTGCTGGTAGTGGTCGGCGAGCTGCACGCTGCCGTCCTCGATGTTGTGGCGGATCTTGAAGTT
CACCTTGATGCCGTTCTTCTGCTTGTCGGCCATGATATAGACGTTGTGGCTGTTGTAGTTGTACTCCAGCTTGTGCCCCAGGATGTTGCCGTC
CTCCTTGAAGTCGATGCCCTTCAGCTCGATGCGGTTCACCAGGGTGTCGCCCTCGAACTTCACCTCGGCGCGGGTCTTGTAGTTGCCGTCGTC
CTTGAAGAAGATGGTGCGCTCCTGGACGTAGCCTTCGGGCATGGCGGACTTGAAGAAGTCGTGCTGCTTCATGTGGTCGGGGTAGCGGCTGAA
GCACTGCACGCATCTGCGGCCAGGGTGGTCACGAGGGTGGGCCAGGGCACGGGCAGCTTGCCGGTGGTGCAGATGAACTTCAGGGTCAGCTTG
CCGTAGGTGGCATCGCCCTCGCCCTCGCCGGACACGCTGAACTTGTGGCCGTTTACGTCGCCGTCCAGCTCGACCAGGATGGGCACCACCCCG
GTGAACAGCTCCTCGCCCTTGCTCACCATGGTGGCTCTCGAGAGCGGATCTGACGGTTCACTAAACCAGCTCTGCTTATATAGACCTCCCACC
GTACACGCCTACCGCCCATTTGCGTCAATGGGGCGGAGTTGTTACGACATTTTGGAAAGTCCCGTTGATTTTGGTGCCAAAACAAACTCCCAT
TGACGTCAATGGGGTGGAGACTTGGAAATCCCCGTGAGTCAAACCGCTATCCACGCCCATTGATGTACTGCCAAAACCGCATCACCATGGTAA
TAGCGATGACTAATACGTAGATGTACTGCCAAGTAGGAAAGTCCCATAAGGTCATGTACTGGGCATAATGCCAGGCGGGCCATTTACCGTCAT
TGACGTCAATAGGGGGCGTACTTGGCATATGATACACTTGATGTACTGCCAAGTGGGCAGTTTACCGTAAATACTCCACCCATTGACGTCAAT
GGAAAGTCCCTATTGGCGTTACTATGGGAACATACGTCATTATTGACGTCAATGGGCGGGGGTCGTTGGGCGGTCAGCCAGGCGGGCCATTTA
CCGTAAGTTATGTAACGCGGAACTCCATATATGGGCTATGAACTAATGACCCCGTAATTGATTACTATTAATAACTAACGCGTGATCTCGAGG
AATTCGAGCTCGGTACCTCGCGAATGCATCTAGATATCACGCGT

### SpCas9-TLRsgRNA

GACTCTTCGCGATGTACGGGCCAGATATACGCGTTGTACAAAAAAGCAGGCTTTAAAGGAACCAATTCAGTCGACTGGATCCGGTACCAAGGT
CGGGCAGGAAGAGGGCCTATTTCCCATGATTCCTTCATATTTGCATATACGATACAAGGCTGTTAGAGAGATAATTAGAATTAATTTGACTGT
AAACACAAAGATATTAGTACAAAATACGTGACGTAGAAAGTAATAATTTCTTGGGTAGTTTGCAGTTTTAAAATTATGTTTTAAAATGGACTA
TCATATGCTTACCGTAACTTGAAAGTATTTCGATTTCTTGGCTTTATATATCTTGTGGAAAGGACGAAACACCGGACTGCACGCATCTGCGGC
CAGTTTTAGAGCTAGAAATAGCAAGTTAAAATAAGGCTAGTCCGTTATCAACTTGAAAAAGTGGCACCGAGTCGGTGCTTTTTTTCTAGACCC
AGCTTTCTTGTACAAAGTTGGCATTAACGCGTTGACATTGATTATTGACTAGTTATTAATAGTAATCAATTACGGGGTCATTAGTTCATAGCC
CATATATGGAGTTCCGCGTTACATAACTTACGGTAAATGGCCCGCCTGGCTGACCGCCCAACGACCCCCGCCCATTGACGTCAATAATGACGT
ATGTTCCCATAGTAACGCCAATAGGGACTTTCCATTGACGTCAATGGGTGGACTATTTACGGTAAACTGCCCACTTGGCAGTACATCAAGTGT
ATCATATGCCAAGTACGCCCCCTATTGACGTCAATGACGGTAAATGGCCCGCCTGGCATTATGCCCAGTACATGACCTTATGGGACTTTCCTA
CTTGGCAGTACATCTACGTATTAGTCATCGCTATTACCATGGTGATGCGGTTTTGGCAGTACATCAATGGGCGTGGATAGCGGTTTGACTCAC
GGGGATTTCCAAGTCTCCACCCCATTGACGTCAATGGGAGTTTGTTTTGGCACCAAAATCAACGGGACTTTCCAAAATGTCGTAACAACTCCG
CCCCATTGACGCAAATGGGCGGTAGGCGTGTACGGTGGGAGGTCTATATAAGCAGAGCTCTCTGGCTAACTAGAGAACCCACTGCTTACTGGC
TTATCGAAATTAATACGACTCACTATAGGGAGACCCAAGCTGGCTAGCGTTTAAACTTAAGCTGATCCACTAGTCCAGTGTGGTGGAATTCGC
CATGGCTCCTAAGAAAAAGCGGAAGGTGGACAAGAAATACTCAATCGGGCTGGACATCGGAACTAACTCAGTGGGGTGGGCAGTCATTACTGA
CGAGTACAAAGTGCCAAGCAAGAAATTTAAGGTCCTGGGCAACACCGATAGGCACTCCATCAAGAAAAATCTGATTGGGGCCCTGCTGTTCGA
CTCTGGAGAGACAGCTGAAGCAACTAGACTGAAAAGGACTGCTAGAAGGCGCTATACCCGGCGAAAGAATCGCATCTGCTACCTGCAGGAGAT
TTTCTCTAACGAAATGGCCAAGGTGGACGATAGTTTCTTTCATCGGCTGGAGGAATCATTCCTGGTCGAGGAAGATAAGAAACACGAGAGACA
TCCTATCTTTGGAAACATTGTGGACGAGGTCGCTTATCACGAAAAATACCCCACCATCTATCATCTGCGCAAGAAACTGGTGGACTCTACAGA
TAAAGCAGACCTGCGGCTGATCTATCTGGCCCTGGCTCACATGATTAAGTTCAGAGGCCATTTTCTGATCGAGGGAGATCTGAACCCAGACAA
TAGCGATGTGGACAAGCTGTTCATCCAGCTGGTCCAGACATACAATCAGCTGTTTGAGGAAAACCCTATTAATGCATCTGGCGTGGACGCAAA
AGCCATCCTGAGTGCCAGGCTGTCTAAGAGTAGAAGGCTGGAGAACCTGATCGCTCAGCTGCCAGGCGAAAAGAAAAACGGCCTGTTTGGAAA
TCTGATTGCACTGTCACTGGGACTGACACCTAACTTCAAGAGCAATTTTGATCTGGCCGAGGACGCTAAACTGCAGCTGAGCAAGGACACTTA
TGACGATGACCTGGATAACCTGCTGGCTCAGATCGGAGATCAGTACGCAGACCTGTTCCTGGCCGCTAAGAATCTGTCTGACGCTATCCTGCT
GAGTGATATTCTGCGGGTGAACACCGAGATTACAAAAGCCCCTCTGTCAGCTAGCATGATCAAGAGATATGACGAGCACCATCAGGATCTGAC
CCTGCTGAAGGCACTGGTGCGCCAGCAGCTGCCCGAGAAGTACAAGGAAATCTTCTTTGATCAGAGTAAGAACGGGTACGCCGGTTATATTGA
CGGCGGAGCTTCACAGGAGGAATTCTACAAGTTTATCAAACCTATTCTGGAGAAGATGGACGGCACCGAGGAACTGCTGGTGAAACTGAATCG
CGAGGACCTGCTGCGCAAGCAGCGGACATTTGATAACGGCTCCATCCCCCACCAGATTCATCTGGGAGAGCTGCACGCAATCCTGCGACGACA
GGAAGACTTCTACCCATTTCTGAAGGATAACCGCGAGAAGATCGAAAAAATTCTGACCTTCCGGATCCCTTACTATGTGGGCCCCCTGGCAAG
GGGTAATTCCCGCTTTGCCTGGATGACACGGAAATCTGAGGAAACAATCACTCCTTGGAACTTCGAGGAAGTGGTCGATAAGGGAGCTTCCGC
ACAGTCTTTCATCGAGAGAATGACAAACTTCGACAAAAACCTGCCAAATGAGAAAGTGCTGCCTAAGCACAGTCTGCTGTACGAGTATTTCAC
AGTCTATAACGAACTGACTAAGGTGAAATACGTCACCGAGGGGATGAGGAAGCCCGCCTTCCTGAGCGGTGAACAGAAGAAAGCTATCGTGGA
CCTGCTGTTTAAAACCAATCGCAAGGTGACAGTCAAGCAGCTGAAGGAGGACTACTTCAAGAAAATTGAATGTTTCGATTCTGTGGAGATCAG
TGGCGTCGAAGACAGATTTAACGCTTCTCTGGGAACCTACCACGATCTGCTGAAGATCATTAAGGATAAAGACTTCCTGGACAACGAGGAAAA
TGAGGATATCCTGGAAGACATTGTGCTGACCCTGACACTGTTTGAGGATCGCGAAATGATCGAGGAACGGCTGAAAACTTATGCCCATCTGTT
CGATGACAAGGTGATGAAACAGCTGAAGCGAAGAAGGTACACCGGCTGGGGACGACTGAGCAGAAAGCTGATCAACGGCATTCGGGACAAACA
GAGTGGAAAGACTATCCTGGACTTTCTGAAATCAGATGGCTTCGCTAACAGAAATTTTATGCAGCTGATTCACGATGACAGCCTGACCTTCAA
AGAGGATATCCAGAAGGCACAGGTGTCCGGGCAGGGTGACTCTCTGCACGAGCATATCGCAAACCTGGCCGGGTCCCCCGCCATCAAGAAAGG
TATTCTGCAGACCGTGAAGGTGGTCGATGAGCTGGTGAAAGTCATGGGCAGGCATAAGCCAGAAAACATCGTGATTGAGATGGCCCGCGAAAA
TCAGACCACACAGAAAGGACAGAAGAACAGCCGCGAGCGGATGAAAAGGATCGAGGAAGGCATTAAGGAACTGGGATCCCAGATCCTGAAAGA
GCACCCTGTGGAAAACACTCAGCTGCAGAATGAGAAGCTGTATCTGTACTATCTGCAGAATGGGCGGGATATGTACGTGGACCAGGAGCTGGA
TATTAACCGACTGTCTGATTACGACGTGGATCATATCGTCCCACAGTCATTCCTGAAAGATGACAGCATTGACAATAAGGTGCTGACCCGGAG
TGACAAAAACCGAGGAAAGAGTGATAATGTCCCTTCAGAGGAAGTGGTCAAGAAAATGAAGAACTACTGGAGACAGCTGCTGAATGCCAAACT
GATCACACAGCGAAAGTTTGATAACCTGACTAAAGCTGAGAGAGGGGGTCTGTCAGAACTGGACAAAGCAGGCTTCATCAAGCGACAGCTGGT
GGAGACCAGACAGATCACAAAGCACGTCGCTCAGATTCTGGATAGCAGGATGAACACAAAGTACGATGAGAATGACAAACTGATCCGCGAAGT
GAAGGTCATTACTCTGAAGTCAAAACTTGTGAGCGACTTCAGAAAGGATTTCCAGTTCTACAAAGTCAGGGAGATCAACAATTATCACCATGC
TCATGACGCATACCTGAACGCAGTGGTCGGGACCGCCCTGATTAAGAAATACCCCAAACTGGAGAGCGAATTCGTGTACGGTGACTATAAGGT
GTACGATGTCAGAAAAATGATCGCCAAGAGTGAGCAGGAAATTGGAAAAGCCACCGCTAAGTATTTCTTTTACTCAAACATCATGAATTTCTT
TAAGACTGAGATCACCCTGGCAAATGGGGAAATCCGAAAGAGACCACTGATTGAGACTAACGGCGAGACCGGAGAAATCGTGTGGGACAAGGG
TAGGGATTTTGCCACAGTGCGCAAGGTCCTGTCCATGCCTCAAGTGAATATTGTCAAGAAAACAGAGGTGCAGACTGGCGGATTCAGTAAGGA
ATCAATTCTGCCCAAACGGAACTCTGATAAGCTGATCGCCCGAAAGAAAGACTGGGATCCCAAGAAATATGGGGGTTTCGACTCCCCAACAGT
GGCTTACTCTGTCCTGGTGGTCGCAAAGGTGGAGAAGGGGAAAAGCAAGAAACTGAAATCCGTCAAGGAGCTGCTGGGTATCACTATTATGGA
GAGGAGCTCCTTCGAGAAGAACCCCATCGATTTTCTGGAGGCTAAAGGCTATAAGGAAGTGAAGAAAGACCTGATCATTAAACTGCCAAAGTA
CAGCCTGTTTGAGCTGGAAAACGGAAGGAAGCGAATGCTGGCATCCGCAGGAGAGCTGCAGAAGGGTAATGAACTGGCCCTGCCTTCTAAGTA
CGTGAACTTCCTGTATCTGGCTAGCCACTACGAGAAGCTGAAAGGCTCCCCCGAGGATAACGAACAGAAACAGCTGTTTGTGGAGCAGCACAA
GCATTATCTGGACGAGATCATTGAACAGATTAGCGAGTTCTCCAAAAGAGTGATCCTGGCTGACGCAAATCTGGATAAGGTCCTGAGCGCATA
CAACAAACACAGAGATAAGCCAATCAGGGAGCAGGCCGAAAATATCATTCATCTGTTCACTCTGACCAACCTGGGAGCCCCTGCAGCCTTCAA

GTATTTTGACACTACCATCGATCGGAAACGATACACATCCACTAAGGAGGTGCTGGACGCTACCCTGATTCACCAGAGCATTACCGGCCTGTA
TGAAACAAGGATTGACCTGTCTCAGCTGGGGGGCGACCTCGAGAGATCTGGCAGCGGAGAGGGCAGAGGAAGTCTTCTAACATGCGGTGACGT
GGAGGAGAATCCCGGCCCTAGGCTCGAAATGACCGAGTACAAGCCCACGGTGCGCCTCGCCACCCGCGACGACGTCCCCCGGGCCGTACGCAC
CCTCGCCGCCGCGTTCGCCGACTACCCCGCCACGCGCCACACCGTCGACCCGGACCGCCACATCGAGCGGGTCACCGAGCTGCAAGAACTCTT
CCTCACGCGCGTCGGGCTCGACATCGGCAAGGTGTGGGTCGCGGACGACGGCGCCGCGGTGGCGGTCTGGACCACGCCGGAGAGCGTCGAAGC
GGGGGCGGTGTTCGCCGAGATCGGCCCGCGCATGGCCGAGTTGAGCGGTTCCCGGCTGGCCGCGCAGCAACAGATGGAAGGCCTCCTGGCGCC
GCACCGGCCCAAGGAGCCCGCGTGGTTCCTGGCCACCGTCGGCGTCTCGCCCGACCACCAGGGCAAGGGTCTGGGCAGCGCCGTCGTGCTCCC
CGGAGTGGAGGCGGCCGAGCGCGCCGGGGTGCCCGCCTTCCTGGAGACCTCCGCGCCCCGCAACCTCCCCTTCTACGAGCGGCTCGGCTTCAC
CGTCACCGCCGACGTCGAGGTGCCCGAAGGACCGCGCACCTGGTGCATGACCCGCAAGCCCGGTGCCTGACGCCCGCCCCACTACCCGCAGCG
CCCGACCGAAAGGAGCGCACGACCCCATGGCTCCGACCGAAGCCACCCGGGGCGGCCCCGCCGACCCCGCACCCGCCCCCGAGGCCCACCGAC
TCTAGAGGGCCCGTTTAAACCCGCTGATCAGCCTCGACTGTGCCTTCTAGTTGCCAGCCATCTGTTGTTTGCCCCTCCCCCGTGCCTTCCTTG
ACCCTGGAAGGTGCCACTCCCACTGTCCTTTCCTAATAAAATGAGGAAATTGCATCGCATTGTCTGAGTAGGTGTCATTCTATTCTGGGGGGT
GGGGTGGGGCAGGACAGCAAGGGGGAGGATTGGGAAGACAATAGCAGGCATGCTGGGGATGCGGTGGGCTCTATGGCTTCTACTGGGCGGTTT
TATGGACAGCAAGCGAACCGGAATTGCCAGCTGGGGCGCCCTCTGGTAAGGTTGGGAAGCCCTGCAAAGTAAACTGGATGGCTTTCTCGCCGC
CAAGGATCTGATGGCGCAGGGGATCAAGCTCTGATCAAGAGACAGGATGAGGATCGTTTCGCATGATTGAACAAGATGGATTGCACGCAGGTT
CTCCGGCCGCTTGGGTGGAGAGGCTATTCGGCTATGACTGGGCACAACAGACAATCGGCTGCTCTGATGCCGCCGTGTTCCGGCTGTCAGCGC
AGGGGCGCCCGGTTCTTTTTGTCAAGACCGACCTGTCCGGTGCCCTGAATGAACTGCAAGACGAGGCAGCGCGGCTATCGTGGCTGGCCACGA
CGGGCGTTCCTTGCGCAGCTGTGCTCGACGTTGTCACTGAAGCGGGAAGGGACTGGCTGCTATTGGGCGAAGTGCCGGGGCAGGATCTCCTGT
CATCTCACCTTGCTCCTGCCGAGAAAGTATCCATCATGGCTGATGCAATGCGGCGGCTGCATACGCTTGATCCGGCTACCTGCCCATTCGACC
ACCAAGCGAAACATCGCATCGAGCGAGCACGTACTCGGATGGAAGCCGGTCTTGTCGATCAGGATGATCTGGACGAAGAGCATCAGGGGCTCG
CGCCAGCCGAACTGTTCGCCAGGCTCAAGGCGAGCATGCCCGACGGCGAGGATCTCGTCGTGACCCATGGCGATGCCTGCTTGCCGAATATCA
TGGTGGAAAATGGCCGCTTTTCTGGATTCATCGACTGTGGCCGGCTGGGTGTGGCGGACCGCTATCAGGACATAGCGTTGGCTACCCGTGATA
TTGCTGAAGAGCTTGGCGGCGAATGGGCTGACCGCTTCCTCGTGCTTTACGGTATCGCCGCTCCCGATTCGCAGCGCATCGCCTTCTATCGCC
TTCTTGACGAGTTCTTCTGAATTATTAACGCTTACAATTTCCTGATGCGGTATTTTCTCCTTACGCATCTGTGCGGTATTTCACACCGCATAC
AGGTGGCACTTTTCGGGGAAATGTGCGCGGAACCCCTATTTGTTTATTTTTCTAAATACATTCAAATATGTATCCGCTCATGAGACAATAACC
CTGATAAATGCTTCAATAATAGCACGTGCTAAAACTTCATTTTTAATTTAAAAGGATCTAGGTGAAGATCCTTTTTGATAATCTCATGACCAA
AATCCCTTAACGTGAGTTTTCGTTCCACTGAGCGTCAGACCCCGTAGAAAAGATCAAAGGATCTTCTTGAGATCCTTTTTTTCTGCGCGTAAT
CTGCTGCTTGCAAACAAAAAAACCACCGCTACCAGCGGTGGTTTGTTTGCCGGATCAAGAGCTACCAACTCTTTTTCCGAAGGTAACTGGCTT
CAGCAGAGCGCAGATACCAAATACTGTCCTTCTAGTGTAGCCGTAGTTAGGCCACCACTTCAAGAACTCTGTAGCACCGCCTACATACCTCGC
TCTGCTAATCCTGTTACCAGTGGCTGCTGCCAGTGGCGATAAGTCGTGTCTTACCGGGTTGGACTCAAGACGATAGTTACCGGATAAGGCGCA
GCGGTCGGGCTGAACGGGGGGTTCGTGCACACAGCCCAGCTTGGAGCGAACGACCTACACCGAACTGAGATACCTACAGCGTGAGCTATGAGA
AAGCGCCACGCTTCCCGAAGGGAGAAAGGCGGACAGGTATCCGGTAAGCGGCAGGGTCGGAACAGGAGAGCGCACGAGGGAGCTTCCAGGGGG
AAACGCCTGGTATCTTTATAGTCCTGTCGGGTTTCGCCACCTCTGACTTGAGCGTCGATTTTTGTGATGCTCGTCAGGGGGGCGGAGCCTATG
GAAAAACGCCAGCAACGCGGCCTTTTTACGGTTCCTGGGCTTTTGCTGGCCTTTTGCTCACATGTTCTT

### TLR-GFP-Donor

GGGGAATTGTGAGCGGATAACAATTCCCCTGTAGAAATAATTTTGTTTAACTTTAATAAGGAGATATACCATGGGCAGCAGCCATCACCATCA
TCACCACAGCCAGGATCCGAGCAAGGGCGAGGAGCTGTTCACCGGGGTGGTGCCCATCCTGGTCGAGCTGGACGGCGACGTAAACGGCCACAA
GTTCAGCGTGTCCGGCGAGGGCGAGGGCGATGCCACCTACGGCAAGCTGACCCTGAAGTTCATCTGCACCACCGGCAAGCTGCCCGTGCCCTG
GCCCACCCTCGTGACCACCCTGACCTACGGCGTGCAGTGCTTCAGCCGCTACCCCGACCACATGAAGCAGCACGACTTCTTCAAGTCCGCCAT
GCCCGAAGGCTACGTCCAGGAGCGCACCATCTTCTTCAAGGACGACGGCAACTACAAGACCCGCGCCGAGGTGAAGTTCGAGGGCGACACCCT
GGTGAACCGCATCGAGCTGAAGGGCATCGACTTCAAGGAGGACGGCAACATCCTGGGGCACAAGCTGGAGTACAACTACAACAGCCACAACGT
CTATATCATGGCCGACAAGCAGAAGAACGGCATCAAGGTGAACTTCAAGATCCGCCACAACATCGAGGACGGCAGCGTGCAGCTCGCCGACCA
CTACCAGCAGAACACCCCCATCGGCGACGGCCCCGTGCTGCTGCCCGACAACCACTACCTGAGCACCCAGTCCGCCCTGAGCAAAGACCCCAA
CGAGAAGCGCGATCACATGGTCCTGCTGGAGTTCGTGACCGCCGCCGGGATCACTCTCGGCATGGACGAGCTGTACAAGTAAGCGGCCGCATA
ATGCTTAAGTCGAACAGAAAGTAATCGTATTGTACACGGCCGCATAATCGAAATTAATACGACTCACTATAGGGGAATTGTGAGCGGATAACA
ATTCCCCATCTTAGTATATTAGTTAAGTATAAGAAGGAGATATACATATGGCAGATCTCAATTGGATATCGGCCGGCCACGCGATCGCTGACG
TCGGTACCCTCGAGTCTGGTAAAGAAACCGCTGCTGCGAAATTTGAACGCCAGCACATGGACTCGTCTACTAGCGCAGCTTAATTAACCTAGG
CTGCTGCCACCGCTGAGCAATAACTAGCATAACCCCTTGGGGCCTCTAAACGGGTCTTGAGGGGTTTTTTGCTGAAACCTCAGGCATTTGAGA
AGCACACGGTCACACTGCTTCCGGTAGTCAATAAACCGGTAAACCAGCAATAGACATAAGCGGCTATTTAACGACCCTGCCCTGAACCGACGA
CCGGGTCGAATTTGCTTTCGAATTTCTGCCATTCATCCGCTTATTATCACTTATTCAGGCGTAGCACCAGGCGTTTAAGGGCACCAATAACTG
CCTTAAAAAAATTACGCCCCGCCCTGCCACTCATCGCAGTACTGTTGTAATTCATTAAGCATTCTGCCGACATGGAAGCCATCACAGACGGCA
TGATGAACCTGAATCGCCAGCGGCATCAGCACCTTGTCGCCTTGCGTATAATATTTGCCCATAGTGAAAACGGGGGCGAAGAAGTTGTCCATA
TTGGCCACGTTTAAATCAAAACTGGTGAAACTCACCCAGGGATTGGCTGAGACGAAAAACATATTCTCAATAAACCCTTTAGGGAAATAGGCC
AGGTTTTCACCGTAACACGCCACATCTTGCGAATATATGTGTAGAAACTGCCGGAAATCGTCGTGGTATTCACTCCAGAGCGATGAAAACGTT
TCAGTTTGCTCATGGAAAACGGTGTAACAAGGGTGAACACTATCCCATATCACCAGCTCACCGTCTTTCATTGCCATACGGAACTCCGGATGA
GCATTCATCAGGCGGGCAAGAATGTGAATAAAGGCCGGATAAAACTTGTGCTTATTTTTCTTTACGGTCTTTAAAAAGGCCGTAATATCCAGC
TGAACGGTCTGGTTATAGGTACATTGAGCAACTGACTGAAATGCCTCAAAATGTTCTTTACGATGCCATTGGGATATATCAACGGTGGTATAT
CCAGTGATTTTTTTCTCCATTTTAGCTTCCTTAGCTCCTGAAAATCTCGATAACTCAAAAAATACGCCCGGTAGTGATCTTATTTCATTATGG
TGAAAGTTGGAACCTCTTACGTGCCGATCAACGTCTCATTTTCGCCAAAAGTTGGCCCAGGGCTTCCCGGTATCAACAGGGACACCAGGATTT
ATTTATTCTGCGAAGTGATCTTCCGTCACAGGTATTTATTCGGCGCAAAGTGCGTCGGGTGATGCTGCCAACTTACTGATTTAGTGTATGATG
GTGTTTTTGAGGTGCTCCAGTGGCTTCTGTTTCTATCAGCTGTCCCTCCTGTTCAGCTACTGACGGGGTGGTGCGTAACGGCAAAAGCACCGC
CGGACATCAGCGCTAGCGGAGTGTATACTGGCTTACTATGTTGGCACTGATGAGGGTGTCAGTGAAGTGCTTCATGTGGCAGGAGAAAAAAGG
CTGCACCGGTGCGTCAGCAGAATATGTGATACAGGATATATTCCGCTTCCTCGCTCACTGACTCGCTACGCTCGGTCGTTCGACTGCGGCGAG
CGGAAATGGCTTACGAACGGGGCGGAGATTTCCTGGAAGATGCCAGGAAGATACTTAACAGGGAAGTGAGAGGGCCGCGGCAAAGCCGTTTTT
CCATAGGCTCCGCCCCCCTGACAAGCATCACGAAATCTGACGCTCAAATCAGTGGTGGCGAAACCCGACAGGACTATAAAGATACCAGGCGTT
TCCCCTGGCGGCTCCCTCGTGCGCTCTCCTGTTCCTGCCTTTCGGTTTACCGGTGTCATTCCGCTGTTATGGCCGCGTTTGTCTCATTCCACG
CCTGACACTCAGTTCCGGGTAGGCAGTTCGCTCCAAGCTGGACTGTATGCACGAACCCCCCGTTCAGTCCGACCGCTGCGCCTTATCCGGTAA
CTATCGTCTTGAGTCCAACCCGGAAAGACATGCAAAAGCACCACTGGCAGCAGCCACTGGTAATTGATTTAGAGGAGTTAGTCTTGAAGTCAT
GCGCCGGTTAAGGCTAAACTGAAAGGACAAGTTTTGGTGACTGCGCTCCTCCAAGCCAGTTACCTCGGTTCAAAGAGTTGGTAGCTCAGAGAA
CCTTCGAAAAACCGCCCTGCAAGGCGGTTTTTTCGTTTTCAGAGCAAGAGATTACGCGCAGACCAAAACGATCTCAAGAAGATCATCTTATTA
ATCAGATAAAATATTTCTAGATTTCAGTGCAATTTATCTCTTCAAATGTAGCACCTGAAGTCAGCCCCATACGATATAAGTTGTAATTCTCAT

GTTAGTCATGCCCCGCGCCCACCGGAAGGAGCTGACTGGGTTGAAGGCTCTCAAGGGCATCGGTCGAGATCCCGGTGCCTAATGAGTGAGCTA
ACTTACATTAATTGCGTTGCGCTCACTGCCCGCTTTCCAGTCGGGAAACCTGTCGTGCCAGCTGCATTAATGAATCGGCCAACGCGCGGGGAG
AGGCGGTTTGCGTATTGGGCGCCAGGGTGGTTTTTCTTTTCACCAGTGAGACGGGCAACAGCTGATTGCCCTTCACCGCCTGGCCCTGAGAGA
GTTGCAGCAAGCGGTCCACGCTGGTTTGCCCCAGCAGGCGAAAATCCTGTTTGATGGTGGTTAACGGCGGGATATAACATGAGCTGTCTTCGG
TATCGTCGTATCCCACTACCGAGATGTCCGCACCAACGCGCAGCCCGGACTCGGTAATGGCGCGCATTGCGCCCAGCGCCATCTGATCGTTGG
CAACCAGCATCGCAGTGGGAACGATGCCCTCATTCAGCATTTGCATGGTTTGTTGAAAACCGGACATGGCACTCCAGTCGCCTTCCCGTTCCG
CTATCGGCTGAATTTGATTGCGAGTGAGATATTTATGCCAGCCAGCCAGACGCAGACGCGCCGAGACAGAACTTAATGGGCCCGCTAACAGCG
CGATTTGCTGGTGACCCAATGCGACCAGATGCTCCACGCCCAGTCGCGTACCGTCTTCATGGGAGAAAATAATACTGTTGATGGGTGTCTGGT
CAGAGACATCAAGAAATAACGCCGGAACATTAGTGCAGGCAGCTTCCACAGCAATGGCATCCTGGTCATCCAGCGGATAGTTAATGATCAGCC
CACTGACGCGTTGCGCGAGAAGATTGTGCACCGCCGCTTTACAGGCTTCGACGCCGCTTCGTTCTACCATCGACACCACCACGCTGGCACCCA
GTTGATCGGCGCGAGATTTAATCGCCGCGACAATTTGCGACGGCGCGTGCAGGGCCAGACTGGAGGTGGCAACGCCAATCAGCAACGACTGTT
TGCCCGCCAGTTGTTGTGCCACGCGGTTGGGAATGTAATTCAGCTCCGCCATCGCCGCTTCCACTTTTTCCCGCGTTTTCGCAGAAACGTGGC
TGGCCTGGTTCACCACGCGGGAAACGGTCTGATAAGAGACACCGGCATACTCTGCGACATCGTATAACGTTACTGGTTTCACATTCACCACCC
TGAATTGACTCTCTTCCGGGCGCTATCATGCCATACCGCGAAAGGTTTTGCGCCATTCGATGGTGTCCGGGATCTCGACGCTCTCCCTTATGC
GACTCCTGCATTAGGAAATTAATACGACTCACTATA

### HBEGF-HDR-Donor-plasmid

GAGGGAGAAGCCAGTAGGAAACATGGCAGAGTGGGCTTCCAGGGCAGAGTAGAGCTCCTGTGGGAAGGTAGGAAGTGCATTTGGATGCATGAT
GTATAGGTATGTGTGTATTTGGGTTTATGTGCATGTAAGTGTGCAAATGTGGATTGACTGTGAGGCATGGCAGGACTGTACAGAGAGGGATCA
TCATGGCGGCAGGTTGAGGCCTCTCTTTCTTCTTCCTTATCCCAGCAAGGACGAGGAGGTGGGAGACATGGAGAGTACTGGCCTTTGGCCACG
TTGTGAGAGAACAATTCCTTTGTGCAGGGTTCACAGGAAATGGAACCTGACCCATTAGGCATCAGCCCCCGCAATTGGTCAGGCAACATCACC
CCTTCCCTGGGTAGGTGTGTGGGTGGAGGGGCTGTGGGTTCCTTAGCCTCTCTCCTAAGCCAAACCCAGCAAACGGCTGCCTTGGCAACCCCT
CAGGGATGACAGCACTGCCATGCTCTCTGGCAGGCATAATGTTGCCACTGTGCCTGAGGCCAACACCCTGCGTCAGGCTGCAAACATCCATTC
CCTTCCCTGTGGGGAGGGAGGCTCTGGGGGCCTTAGTGGGAGACTCTGGACAGGGCCAAGAGACTGTTGTATGCACACTGCCTCCAGCCTGTC
AAGAAGGCGGCGTGCCTGGCATCCCTTCTACTGGTGATTGGTGCAGATCCCTTAG

### HDRT-plasmid

CCTGCAGGCAGCTGCGCGCTCGCTCGCTCACTGAGGCCGCCCGGGCGTCGGGCGACCTTTGGTCGCCCGGCCTCAGTGAGCGAGCGAGCGCGC
AGAGAGGGAGTGGCCAACTCCATCACTAGGGGTTCCTGCGGCCGCATCGATTGAATTCCCCGGGGATCCTCTAGAGTCCTCAGCGCTCTTCAG
CTGTGAAATCATGGCCTCTTGGCCAAGATTGATAGCTTGTGCCTGTCCCTGAGTCCCAGTCCATCACGAGCAGCTGGTTTCTAAGATGCTATT
TCCCGTATAAAGCATGAGACCGTGACTTGCCAGCCCCACAGAGCCCCGCCCTTGTCCATCACTGGCATCTGGACTCCAGCCTGGGTTGGGGCA
AAGAGGGAAATGAGATCATGTCCTAACCCTGATCCTCTTGTAAGCTTCTGACCTCTTCTCTTCCTCCCACAGGGCCTCGAGCTGGCAGCGGAG
AGGGCAGAGGAAGTCTTCTAACATGCGGTGACGTGGAGGAGAATCCCGGCCCTAGGCTCGAGATGGTGAGCAAGGGCGAGGAGCTGTTCACCG
GGGTGGTGCCCATCCTGGTCGAGCTGGACGGCGACGTAAACGGCCACAAGTTCAGCGTGTCCGGCGAGGGCGAGGGCGATGCCACCTACGGCA
AGCTGACCCTGAAGTTCATCTGCACCACCGGCAAGCTGCCCGTGCCCTGGCCCACCCTCGTGACCACCTTCACCTACGGCGTGCAGTGCTTCG
CCCGCTACCCCGACCACATGAAGCAGCACGACTTCTTCAAGTCCGCCATGCCCGAAGGCTACGTCCAGGAGCGCACCATCTTCTTCAAGGACG
ACGGCAACTACAAGACCCGCGCCGAGGTGAAGTTCGAGGGCGACACCCTGGTGAACCGCATCGAGCTGAAGGGCATCGACTTCAAGGAGGACG
GCAACATCCTGGGGCACAAGCTGGAGTACAACTACAACAGCCACAAGGTCTATATCACCGCCGACAAGCAGAAGAACGGCATCAAGGTGAACT
TCAAGACCCGCCACAACATCGAGGACGGCAGCGTGCAGCTCGCCGACCACTACCAGCAGAACACCCCCATCGGCGACGGCCCCGTGCTGCTGC
CCGACAACCACTACCTGAGCACCCAGTCCGCCCTGAGCAAAGACCCCAACGAGAAGCGCGATCACATGGTCCTGCTGGAGTTCGTGACCGCCG
CCGGGATCACTCTCGGCATGGACGAGCTGTACAAGTAATCTAGAGGGCCCGTTTAAACCCGCTGATCAGCCTCGATAAGATACATTGATGAGT
TTGGACAAACCACAACTAGAATGCAGTGAAAAAAATGCTTTATTTGTGAAATTTGTGATGCTATTGCTTTATTTGTAACCATTATAAGCTGCA
ATAAACAAGTTGTCGACACCCTGCCGTGTACCAGCTGAGAGACTCTAAATCCAGTGACAAGTCTGTCTGCCTATTCACCGATTTTGATTCTCA
AACAAATGTGTCACAAAGTAAGGATTCTGATGTGTATATCACAGACAAAACTGTGCTAGACATGAGGTCTATGGACTTCAAGAGCAACAGTGC
TGTGGCCTGGAGCAACAAATCTGACTTTGCATGTGCAAACGCCTTCAACAACAGCATTATTCCAGAAGACACCTTCTTCCCGCTCTTCACAGC
CCTCAGCTCTGATTTTGTAGGTAACCACGTGCGGACCGAGCGGCCGCAGGAACCCCTAGTGATGGAGTTGGCCACTCCCTCTCTGCGCGCTCG
CTCGCTCACTGAGGCCGGGCGACCAAAGGTCGCCCGACGCCCGGGCTTTGCCCGGGCGGCCTCAGTGAGCGAGCGAGCGCGCAGCTGCCTGCA
GGGGCGCCTGATGCGGTATTTTCTCCTTACGCATCTGTGCGGTATTTCACACCGCATACGTCAAAGCAACCATAGTACGCGCCCTGTAGCGGC
GCATTAAGCGCGGCGGGTGTGGTGGTTACGCGCAGCGTGACCGCTACACTTGCCAGCGCCCTAGCGCCCGCTCCTTTCGCTTTCTTCCCTTCC
TTTCTCGCCACGTTCGCCGGCTTTCCCCGTCAAGCTCTAAATCGGGGGCTCCCTTTAGGGTTCCGATTTAGTGCTTTACGGCACCTCGACCCC
AAAAAACTTGATTTGGGTGATGGTTCACGTAGTGGGCCATCGCCCTGATAGACGGTTTTTCGCCCTTTGACGTTGGAGTCCACGTTCTTTAAT
AGTGGACTCTTGTTCCAAACTGGAACAACACTCAACCCTATCTCGGGCTATTCTTTTGATTTATAAGGGATTTTGCCGATTTCGGCCTATTGG
TTAAAAAATGAGCTGATTTAACAAAAATTTAACGCGAATTTTAACAAAATATTAACGTTTACAATTTTATGGTGCACTCTCAGTACAATCTGC
TCTGATGCCGCATAGTTAAGCCAGCCCCGACACCCGCCAACACCCGCTGACGCGCCCTGACGGGCTTGTCTGCTCCCGGCATCCGCTTACAGA
CAAGCTGTGACCGTCTCCGGGAGCTGCATGTGTCAGAGGTTTTCACCGTCATCACCGAAACGCGCGAGACGAAAGGGCCTCGTGATACGCCTA
TTTTTATAGGTTAATGTCATGATAATAATGGTTTCTTAGACGTCAGGTGGCACTTTTCGGGGAAATGTGCGCGGAACCCCTATTTGTTTATTT
TTCTAAATACATTCAAATATGTATCCGCTCATGAGACAATAACCCTGATAAATGCTTCAATAATATTGAAAAAGGAAGAGTATGAGTATTCAA
CATTTCCGTGTCGCCCTTATTCCCTTTTTTGCGGCATTTTGCCTTCCTGTTTTTGCTCACCCAGAAACGCTGGTGAAAGTAAAAGATGCTGAA
GATCAGTTGGGTGCACGAGTGGGTTACATCGAACTGGATCTCAACAGCGGTAAGATCCTTGAGAGTTTTCGCCCCGAAGAACGTTTTCCAATG
ATGAGCACTTTTAAAGTTCTGCTATGTGGCGCGGTATTATCCCGTATTGACGCCGGGCAAGAGCAACTCGGTCGCCGCATACACTATTCTCAG
AATGACTTGGTTGAGTACTCACCAGTCACAGAAAAGCATCTTACGGATGGCATGACAGTAAGAGAATTATGCAGTGCTGCCATAACCATGAGT
GATAACACTGCGGCCAACTTACTTCTGACAACGATCGGAGGACCGAAGGAGCTAACCGCTTTTTTGCACAACATGGGGGATCATGTAACTCGC
CTTGATCGTTGGGAACCGGAGCTGAATGAAGCCATACCAAACGACGAGCGTGACACCACGATGCCTGTAGCAATGGCAACAACGTTGCGCAAA
CTATTAACTGGCGAACTACTTACTCTAGCTTCCCGGCAACAATTAATAGACTGGATGGAGGCGGATAAAGTTGCAGGACCACTTCTGCGCTCG
GCCCTTCCGGCTGGCTGGTTTATTGCTGATAAATCTGGAGCCGGTGAGCGTGGGTCTCGCGGTATCATTGCAGCACTGGGGCCAGATGGTAAG
CCCTCCCGTATCGTAGTTATCTACACGACGGGGAGTCAGGCAACTATGGATGAACGAAATAGACAGATCGCTGAGATAGGTGCCTCACTGATT
AAGCATTGGTAACTGTCAGACCAAGTTTACTCATATATACTTTAGATTGATTTAAAACTTCATTTTTAATTTAAAAGGATCTAGGTGAAGATC
CTTTTTGATAATCTCATGACCAAAATCCCTTAACGTGAGTTTTCGTTCCACTGAGCGTCAGACCCCGTAGAAAAGATCAAAGGATCTTCTTGA
GATCCTTTTTTTCTGCGCGTAATCTGCTGCTTGCAAACAAAAAAACCACCGCTACCAGCGGTGGTTTGTTTGCCGGATCAAGAGCTACCAACT
CTTTTTCCGAAGGTAACTGGCTTCAGCAGAGCGCAGATACCAAATACTGTCCTTCTAGTGTAGCCGTAGTTAGGCCACCACTTCAAGAACTCT
GTAGCACCGCCTACATACCTCGCTCTGCTAATCCTGTTACCAGTGGCTGCTGCCAGTGGCGATAAGTCGTGTCTTACCGGGTTGGACTCAAGA
CGATAGTTACCGGATAAGGCGCAGCGGTCGGGCTGAACGGGGGGTTCGTGCACACAGCCCAGCTTGGAGCGAACGACCTACACCGAACTGAGA

TACCTACAGCGTGAGCTATGAGAAAGCGCCACGCTTCCCGAAGGGAGAAAGGCGGACAGGTATCCGGTAAGCGGCAGGGTCGGAACAGGAGAG
CGCACGAGGGAGCTTCCAGGGGGAAACGCCTGGTATCTTTATAGTCCTGTCGGGTTTCGCCACCTCTGACTTGAGCGTCGATTTTTGTGATGC
TCGTCAGGGGGGCGGAGCCTATGGAAAAACGCCAGCAACGCGGCCTTTTTACGGTTCCTGGCCTTTTGCTGGCCTTTTGCTCACATGT

**Supplementary Table S1**. sgRNA target sequences used in this study.

| Figure | Target | Sequence |
|---|---|---|
| Fig. 1; Supp. Fig. 1 | TLR | ACTGCACGCATCTGCGGCCA |
| Fig. 2,3,4; Supp. Fig. 2, 3, 5, 6, 7 | gDel / STAT1 | GAATGAGGGTCCTTTGGGAA |
| Fig. 2,3,4; Supp. Fig. 2, 3, 5, 6, 7 | gIns / CD34 | ATAGGAGAAGATGATGTATA |
| Fig. 2,3,4; Supp. Fig. 2, 3, 5, 6, 7 | gMej / CD34 | TTCATGAGTCTTGACAACAA |
| Fig. 5 | HEK3 | GGCCCAGACTGAGCACGTGA |
| Fig. 5 | HEK4 | GGCACTGCGGCTGGAGGTGG |
| Fig. 5; Supp. Fig. 8 | HBEGF | GGGTGATGTTGCCTGACCGG |
| Fig. 5; Supp. Fig. 8 | PCSK9 | CAGGTTCCACGGGATGCTCT |
| Fig. 6; Supp. Fig. 9, 10 | CTNNB1 | TTTTAAAACTTCTTACCTAA |
| Fig. 6; Supp. Fig. 9, 10 | HDAC2 | GGTGAGACTGTCAAATTCAG |
| Fig. 6; Supp. Fig. 9, 10 | MAPK8 | TTGACAGACGACGATGATGA |
| Fig. 6; Supp. Fig. 9, 10 | NRAS | TGAAATGACTGAGTACAAAC |
| Fig. 6; Supp. Fig. 9, 10 | NR3C1 | TAAGGCAACCATTCTTATTA |
| Fig. 6; Supp. Fig. 9, 10 | TRAC | CAGGGTTCTGGATATCTGT |

**Supplementary Table S2**. Primer and probes used in this study.

| | **<u>NGS- Amplicon Sequencing</u>** |
|---|---|
| **Name** | **Sequence** |
| gMej_fwd_1 | TCGTCGGCAGCGTCAGATGTGTATAAGAGACAGCCTGAAGTCAGGAAATTGCATCAG |
| gMej_fwd_2 | TCGTCGGCAGCGTCAGATGTGTATAAGAGACAGAAGCCCTGAAGTCAGGAAATTGCATCAG |
| gMej_fwd_3 | TCGTCGGCAGCGTCAGATGTGTATAAGAGACAGCGTTCCTGAAGTCAGGAAATTGCATCAG |
| gMej_fwd_4 | TCGTCGGCAGCGTCAGATGTGTATAAGAGACAGGTCACCTGAAGTCAGGAAATTGCATCAG |
| gMej_fwd_5 | TCGTCGGCAGCGTCAGATGTGTATAAGAGACAGACAGCCTGAAGTCAGGAAATTGCATCAG |
| gMej_fwd_6 | TCGTCGGCAGCGTCAGATGTGTATAAGAGACAGCAGTCCTGAAGTCAGGAAATTGCATCAG |
| gMej_fwd_7 | TCGTCGGCAGCGTCAGATGTGTATAAGAGACAGTGTCCCTGAAGTCAGGAAATTGCATCAG |
| gMej_rev_1 | GTCTCGTGGGCTCGGAGATGTGTATAAGAGACAGAGGCTGGTACTTCCAAGGGTA |
| gMej_rev_2 | GTCTCGTGGGCTCGGAGATGTGTATAAGAGACAGAGCAAGGCTGGTACTTCCAAGGGTA |
| gMej_rev_3 | GTCTCGTGGGCTCGGAGATGTGTATAAGAGACAGCCTTAGGCTGGTACTTCCAAGGGTA |
| gMej_rev_4 | GTCTCGTGGGCTCGGAGATGTGTATAAGAGACAGGAACAGGCTGGTACTTCCAAGGGTA |

| | |
|---|---|
| gMej_rev_5 | GTCTCGTGGGCTCGGAGATGTGTATAAGAGACAGTTGGAGGCTGGTACTTCCAAGGGTA |
| gMej_rev_6 | GTCTCGTGGGCTCGGAGATGTGTATAAGAGACAGACAGAGGCTGGTACTTCCAAGGGTA |
| gMej_rev_7 | GTCTCGTGGGCTCGGAGATGTGTATAAGAGACAGCAGTAGGCTGGTACTTCCAAGGGTA |
| gMej_rev_8 | GTCTCGTGGGCTCGGAGATGTGTATAAGAGACAGCTACAGGCTGGTACTTCCAAGGGTA |
| gMej_rev_9 | GTCTCGTGGGCTCGGAGATGTGTATAAGAGACAGGATGAGGCTGGTACTTCCAAGGGTA |
| gMej_rev_10 | GTCTCGTGGGCTCGGAGATGTGTATAAGAGACAGGGATAGGCTGGTACTTCCAAGGGTA |
| gMej_rev_11 | GTCTCGTGGGCTCGGAGATGTGTATAAGAGACAGTGTCAGGCTGGTACTTCCAAGGGTA |
| gIns_fwd_1 | TCGTCGGCAGCGTCAGATGTGTATAAGAGACAGCCAGCCAACGTTTCAACTCC |
| gIns_fwd_2 | TCGTCGGCAGCGTCAGATGTGTATAAGAGACAGAAGCCCAGCCAACGTTTCAACTCC |
| gIns_fwd_3 | TCGTCGGCAGCGTCAGATGTGTATAAGAGACAGCGTTCCAGCCAACGTTTCAACTCC |
| gIns_fwd_4 | TCGTCGGCAGCGTCAGATGTGTATAAGAGACAGGTCACCAGCCAACGTTTCAACTCC |
| gIns_fwd_5 | TCGTCGGCAGCGTCAGATGTGTATAAGAGACAGACAGCCAGCCAACGTTTCAACTCC |
| gIns_fwd_6 | TCGTCGGCAGCGTCAGATGTGTATAAGAGACAGCAGTCCAGCCAACGTTTCAACTCC |
| gIns_fwd_7 | TCGTCGGCAGCGTCAGATGTGTATAAGAGACAGTGTCCCAGCCAACGTTTCAACTCC |
| gIns_rev_1 | GTCTCGTGGGCTCGGAGATGTGTATAAGAGACAGTGAGGGGCAATCTGCCATTT |
| gIns_rev_2 | GTCTCGTGGGCTCGGAGATGTGTATAAGAGACAGAGCATGAGGGGCAATCTGCCATTT |
| gIns_rev_3 | GTCTCGTGGGCTCGGAGATGTGTATAAGAGACAGCCTTTGAGGGGCAATCTGCCATTT |
| gIns_rev_4 | GTCTCGTGGGCTCGGAGATGTGTATAAGAGACAGGAACTGAGGGGCAATCTGCCATTT |
| gIns_rev_5 | GTCTCGTGGGCTCGGAGATGTGTATAAGAGACAGTTGGTGAGGGGCAATCTGCCATTT |
| gIns_rev_6 | GTCTCGTGGGCTCGGAGATGTGTATAAGAGACAGACAGTGAGGGGCAATCTGCCATTT |
| gIns_rev_7 | GTCTCGTGGGCTCGGAGATGTGTATAAGAGACAGCAGTTGAGGGGCAATCTGCCATTT |
| gIns_rev_8 | GTCTCGTGGGCTCGGAGATGTGTATAAGAGACAGCTACTGAGGGGCAATCTGCCATTT |
| gIns_rev_9 | GTCTCGTGGGCTCGGAGATGTGTATAAGAGACAGGATGTGAGGGGCAATCTGCCATTT |
| gIns_rev_10 | GTCTCGTGGGCTCGGAGATGTGTATAAGAGACAGGGATTGAGGGGCAATCTGCCATTT |
| gIns_rev_11 | GTCTCGTGGGCTCGGAGATGTGTATAAGAGACAGTGTCTGAGGGGCAATCTGCCATTT |
| gDel_fwd_1 | TCGTCGGCAGCGTCAGATGTGTATAAGAGACAGGGCATTCTCTGAAGAGTGGGTT |

| | |
|---|---|
| gDel_fwd_2 | TCGTCGGCAGCGTCAGATGTGTATAAGAGACAGAAGCGGCATTCTCTGAAGAG TGGGTT |
| gDel_fwd_3 | TCGTCGGCAGCGTCAGATGTGTATAAGAGACAGCGTTGGCATTCTCTGAAGAG TGGGTT |
| gDel_fwd_4 | TCGTCGGCAGCGTCAGATGTGTATAAGAGACAGGTCAGGCATTCTCTGAAGAG TGGGTT |
| gDel_fwd_5 | TCGTCGGCAGCGTCAGATGTGTATAAGAGACAGACAGGGCATTCTCTGAAGAG TGGGTT |
| gDel_fwd_6 | TCGTCGGCAGCGTCAGATGTGTATAAGAGACAGCAGTGGCATTCTCTGAAGAG TGGGTT |
| gDel_fwd_7 | TCGTCGGCAGCGTCAGATGTGTATAAGAGACAGTGTCGGCATTCTCTGAAGAG TGGGTT |
| gDel_rev_1 | GTCTCGTGGGCTCGGAGATGTGTATAAGAGACAGTTCCTACGTCAAGCAGTTC CC |
| gDel_rev_2 | GTCTCGTGGGCTCGGAGATGTGTATAAGAGACAGAGCATTCCTACGTCAAGCA GTTCCC |
| gDel_rev_3 | GTCTCGTGGGCTCGGAGATGTGTATAAGAGACAGCCTTTTCCTACGTCAAGCA GTTCCC |
| gDel_rev_4 | GTCTCGTGGGCTCGGAGATGTGTATAAGAGACAGGAACTTCCTACGTCAAGCA GTTCCC |
| gDel_rev_5 | GTCTCGTGGGCTCGGAGATGTGTATAAGAGACAGTTGGTTCCTACGTCAAGCA GTTCCC |
| gDel_rev_6 | GTCTCGTGGGCTCGGAGATGTGTATAAGAGACAGACAGTTCCTACGTCAAGCA GTTCCC |
| gDel_rev_7 | GTCTCGTGGGCTCGGAGATGTGTATAAGAGACAGCAGTTTCCTACGTCAAGCA GTTCCC |
| gDel_rev_8 | GTCTCGTGGGCTCGGAGATGTGTATAAGAGACAGCTACTTCCTACGTCAAGCA GTTCCC |
| gDel_rev_9 | GTCTCGTGGGCTCGGAGATGTGTATAAGAGACAGGATGTTCCTACGTCAAGCA GTTCCC |
| gDel_rev_10 | GTCTCGTGGGCTCGGAGATGTGTATAAGAGACAGGGATTTCCTACGTCAAGCA GTTCCC |
| gDel_rev_11 | GTCTCGTGGGCTCGGAGATGTGTATAAGAGACAGTGTCTTCCTACGTCAAGCA GTTCCC |
| HEK3_fwd | TCGTCGGCAGCGTCAGATGTGTATAAGAGACAGCCATGCAATTAGTCTATTTC TGCTGCA |
| HEK3_rev | GTCTCGTGGGCTCGGAGATGTGTATAAGAGACAGCAGCCCAGCCAAACTTGTC AACCAGTA |
| HEK3_OT1_f wd | TCGTCGGCAGCGTCAGATGTGTATAAGAGACAGGGAGAAGCATGAACCAGTCA AAAAGTT |
| HEK3_OT1_r ev | GTCTCGTGGGCTCGGAGATGTGTATAAGAGACAGCGTGGAGCCACTGATGCTC ATGTCATC |
| HEK3_OT2_f wd | TCGTCGGCAGCGTCAGATGTGTATAAGAGACAGGAGACTTAGTGAGACTTGAA ACCTATC |
| HEK3_OT2_r ev | GTCTCGTGGGCTCGGAGATGTGTATAAGAGACAGTGGGAGAGTTAACCTGGGT TCAGTAGA |
| HEK3_OT3_f wd | TCGTCGGCAGCGTCAGATGTGTATAAGAGACAGATTCCTGGTCCAAAGGCCCA AGAACCT |
| HEK3_OT3_r ev | GTCTCGTGGGCTCGGAGATGTGTATAAGAGACAGCAGCCGGCGATGAGTAAGA GTGATGTG |
| HEK3_OT4_f wd | TCGTCGGCAGCGTCAGATGTGTATAAGAGACAGCTCATCTTAATCTGCTCAGC CAGAATT |

| | |
|---|---|
| HEK3_OT4_rev | GTCTCGTGGGCTCGGAGATGTGTATAAGAGACAGACAAAAGCTATGATGTGATGTGACTGG |
| HEK4_fwd | TCGTCGGCAGCGTCAGATGTGTATAAGAGACAGTGGAAGGAAGGGAGGAAGGG |
| HEK4_rev | GTCTCGTGGGCTCGGAGATGTGTATAAGAGACAGCCTTTCAACCCGAACGGAGA |
| HEK4_OT1_fwd | TCGTCGGCAGCGTCAGATGTGTATAAGAGACAGCTGGGGCTGAAGATCCCTAG |
| HEK4_OT1_rev | GTCTCGTGGGCTCGGAGATGTGTATAAGAGACAGCATTCCGGATGATTCTCCTACTT |
| HEK4_OT2_fwd | TCGTCGGCAGCGTCAGATGTGTATAAGAGACAGCATGAGAGCAAGGGAGCC |
| HEK4_OT2_rev | GTCTCGTGGGCTCGGAGATGTGTATAAGAGACAGGTGTGTGTGTTTGTGTGTGTG |
| HEK4_OT3_fwd | TCGTCGGCAGCGTCAGATGTGTATAAGAGACAGGTGAGCTCATTTCCACCAGAACTCAGC |
| HEK4_OT3_rev | GTCTCGTGGGCTCGGAGATGTGTATAAGAGACAGGCAGAGTAGATAAGGACAGATGGGAAA |
| HBEGF_fwd_1 | TCGTCGGCAGCGTCAGATGTGTATAAGAGACAGAAGC |
| HBEGF_fwd_2 | TCGTCGGCAGCGTCAGATGTGTATAAGAGACAGCGTT |
| HBEGF_fwd_3 | TCGTCGGCAGCGTCAGATGTGTATAAGAGACAGGTCA |
| HBEGF_fwd_4 | TCGTCGGCAGCGTCAGATGTGTATAAGAGACAGACAG |
| HBEGF_fwd_5 | TCGTCGGCAGCGTCAGATGTGTATAAGAGACAGCAGT |
| HBEGF_fwd_6 | TCGTCGGCAGCGTCAGATGTGTATAAGAGACAGTGTC |
| HBEGF_rev_1 | GTCTCGTGGGCTCGGAGATGTGTATAAGAGACAGAGCA |
| HBEGF_rev_2 | GTCTCGTGGGCTCGGAGATGTGTATAAGAGACAGCCTT |
| HBEGF_rev_3 | GTCTCGTGGGCTCGGAGATGTGTATAAGAGACAGGAAC |
| HBEGF_rev_4 | GTCTCGTGGGCTCGGAGATGTGTATAAGAGACAGTTGG |
| HBEGF_rev_5 | GTCTCGTGGGCTCGGAGATGTGTATAAGAGACAGACAG |
| HBEGF_rev_6 | GTCTCGTGGGCTCGGAGATGTGTATAAGAGACAGCAGT |
| HBEGF_rev_7 | GTCTCGTGGGCTCGGAGATGTGTATAAGAGACAGCTAC |
| HBEGF_rev_8 | GTCTCGTGGGCTCGGAGATGTGTATAAGAGACAGGATG |
| HBEGF_rev_9 | GTCTCGTGGGCTCGGAGATGTGTATAAGAGACAGGGAT |
| HBEGF_rev_10 | GTCTCGTGGGCTCGGAGATGTGTATAAGAGACAGTGTC |
| PCSK9_fwd_1 | TCGTCGGCAGCGTCAGATGTGTATAAGAGACAGAAGC |

| | |
|---|---|
| PCSK9_fwd_2 | TCGTCGGCAGCGTCAGATGTGTATAAGAGACAGCGTT |
| PCSK9_fwd_3 | TCGTCGGCAGCGTCAGATGTGTATAAGAGACAGGTCA |
| PCSK9_fwd_4 | TCGTCGGCAGCGTCAGATGTGTATAAGAGACAGACAG |
| PCSK9_fwd_5 | TCGTCGGCAGCGTCAGATGTGTATAAGAGACAGCAGT |
| PCSK9_fwd_6 | TCGTCGGCAGCGTCAGATGTGTATAAGAGACAGTGTC |
| PCSK9_rev_1 | GTCTCGTGGGCTCGGAGATGTGTATAAGAGACAGAGCA |
| PCSK9_rev_2 | GTCTCGTGGGCTCGGAGATGTGTATAAGAGACAGCCTT |
| PCSK9_rev_3 | GTCTCGTGGGCTCGGAGATGTGTATAAGAGACAGGAAC |
| PCSK9_rev_4 | GTCTCGTGGGCTCGGAGATGTGTATAAGAGACAGTTGG |
| PCSK9_rev_5 | GTCTCGTGGGCTCGGAGATGTGTATAAGAGACAGACAG |
| PCSK9_rev_6 | GTCTCGTGGGCTCGGAGATGTGTATAAGAGACAGCAGT |
| PCSK9_rev_7 | GTCTCGTGGGCTCGGAGATGTGTATAAGAGACAGCTAC |
| PCSK9_rev_8 | GTCTCGTGGGCTCGGAGATGTGTATAAGAGACAGGATG |
| PCSK9_rev_9 | GTCTCGTGGGCTCGGAGATGTGTATAAGAGACAGGGAT |
| PCSK9_rev_10 | GTCTCGTGGGCTCGGAGATGTGTATAAGAGACAGTGTC |
| CTNNB1_fwd | TCGTCGGCAGCGTCAGATGTGTATAAGAGACAGTTCACTCTGGTGGATATGGCCAGG |
| CTNNB1_rev | GTCTCGTGGGCTCGGAGATGTGTATAAGAGACAGCCTAAACCACTCCCACCCTACCAA |
| HDAC2_fwd | TCGTCGGCAGCGTCAGATGTGTATAAGAGACAGTGTGCCAGCCTTTACAGGATAACT |
| HDAC2_rev | GTCTCGTGGGCTCGGAGATGTGTATAAGAGACAGCCATGCCAAAGTAGTATAAAATGAAGCCA |
| MAPK8_fwd | TCGTCGGCAGCGTCAGATGTGTATAAGAGACAGGGCAGTCAGTGCAGTGCAAGTAG |
| MAPK8_rev | GTCTCGTGGGCTCGGAGATGTGTATAAGAGACAGAATGACTAACCGACTCCCCATCCC |
| NR3C1_fwd | TCGTCGGCAGCGTCAGATGTGTATAAGAGACAGCCCACTGACCAATTTGGAAGCCTG |
| NR3C1_rev | GTCTCGTGGGCTCGGAGATGTGTATAAGAGACAGTGACGACTCAACTGCTTCTGTTGC |
| NRAS_fwd | TCGTCGGCAGCGTCAGATGTGTATAAGAGACAGACTACTCCAGAAGTGTGAGGCCGA |
| NRAS_rev | GTCTCGTGGGCTCGGAGATGTGTATAAGAGACAGCCGACAAGTGAGAGACAGGATCAGG |
| | **TLR Junction PCR** |
| **Name** | **Sequence** |
| 5' junction puro/AVVS1_fwd | cccctatgtccacttcagga |

| | |
|---|---|
| 5' junction puro/AVVS1_rev | tgaggaagagttcttgcagct |
| 3' juntion TLR gfp/AVVS1_fwd | gctcgaccaggatgggcac |
| 3' juntion TLR gfp/AVVS1_rev | acaggaggtgggggttagac |
| | **ddPCR assays** |
| | **TLR cell line** |
| **Name** | **Sequence** |
| Puromycin-fwd | gtcaccgagctgcaagaa |
| Puromycin-rev | caccttgccgatgtcgag |
| Puromycin-Probe | ctcttcctcacgcgcgtcgg |
| | **Translocation** |
| **Name** | **Sequence** |
| PCSK9-HBEGF_1_F | TCGACTACATCGAGGAGGAC |
| PCSK9-HBEGF_1_IN | ACGGCTGCCTTGGCAACCCCTCA |
| PCSK9-HBEGF_1_R | AGGCACAGTGGCAACATTAT |
| | **HiBiT integration** |
| **Name** | **Sequence** |
| CTNNB1_fwd | TTCACTCTGGTGGATATGGCCAGG |
| CTNNB1_rev | CCTAAACCACTCCCACCCTACCAA |
| CTNNB1_Probe 1 | /5HEX/TGGGTGGCC/ZEN/ACCACCCTGGT/3IABkFQ/ |
| HDAC2_fwd | TGTGCCAGCCTTTACAGGATAACT |
| HDAC2_rev | CCATGCCAAAGTAGTATAAAATGAAGCCA |
| HDAC2-Probe 1 | /5HEX/AGCATGAGC/ZEN/CTACAGGACCTGATCA/3IABkFQ/ |
| MAPK8_fwd | GGCAGTCAGTGCAGTGCAAGTAG |
| MAPK8_rev | AATGACTAACCGACTCCCCATCCC |
| MAPK8-Probe 1 | /5HEX/TGGGCCTCT/ZEN/GGGCTGCTGTA/3IABkFQ/ |
| NRAS_fwd | ACTACTCCAGAAGTGTGAGGCCGA |
| NRAS_rev | CCGACAAGTGAGAGACAGGATCAGG |
| NRAS-Probe 1 | /5HEX/AGGCCCAGT/ZEN/GGTAGCCCGCT/3IABkFQ/ |
| NR3C1_fwd | CCCACTGACCAATTTGGAAGCCTG |
| NR3C1_rev | TGACGACTCAACTGCTTCTGTTGC |

| | | |
|---|---|---|
| NR3C1-Probe 1 | /5HEX/ACGCACTAC/ZEN/ATGTGGTTTATAGAGGGCC/3IABkFQ/ | |
| HiBiT-Probe 2 | /56-FAM/TGAGCGGCT/ZEN/GGCGGCTGTTCAAGAAGATTA/3IABkFQ/ | |
| | **HaloTag-HiBiT integration** | |
| | Single Amplicon Method | |
| **Name** | **Sequence** | **Flank** |
| MAPK8_fwd | AACCTCGTGTAAAGTGGGGCAGT | NA |
| MAPK8_rev | ATTTTCCCTCACTCCTCTGGGGTT | NA |
| MAPK8-Probe1 | /5HEX/TGGGCCTCT/ZEN/GGGCTGCTGTA/3IABkFQ/ | NA |
| NRAS_fwd | TTTCCCGGCTGTGGTCCTAAATCT | NA |
| NRAS_rev | CCTGTGTGGTAGGCAGGACAAGTT | NA |
| NRAS-Probe1 | /5HEX/AGGCCCAGT/ZEN/GGTAGCCCGCT/3IABkFQ/ | NA |
| NR3C1_fwd | GCCAGAACTGGCAGCGGTTTTATC | NA |
| NR3C1_rev | AGTGCCATCAGGTTAGAAGCACCA | NA |
| NR3C1-Probe1 | /5HEX/ACGCACTAC/ZEN/ATGTGGTTTATAGAGGGCC/3IABkFQ/ | NA |
| HiBiT | /56-FAM/TGAGCGGCT/ZEN/GGCGGCTGTTCAAGAAGATTA/3IABkFQ/ | NA |
| | 5' Flank and 3' Flank Method | |
| **Name** | **Sequence** | **Flank** |
| HDAC2_fwd | TCCAAGGACAACAGTGGTGAAAAA | 5' |
| HDAC2_rev | CTCGCCCAGGACTTCCACATAATGG | 5' |
| HDAC2-Probe1 | /5HEX/AGCATGAGC/ZEN/CTACAGGACCTGATCA/3IABkFQ/ | 5' |
| HDAC2_fwd | GGCAAGTCTGGTCAATACAAGTTCTGG | 3' |
| HDAC2_rev | CTAACTGCAAGGCTGTGGACATCGG | 3' |
| HiBiT | /56-FAM/TGAGCGGCT/ZEN/GGCGGCTGTTCAAGAAGATTA/3IABkFQ/ | 3' |
| MAPK8_fwd | AACCTCGTGTAAAGTGGGGCAGT | 5' |
| MAPK8_rev | CTCGCCCAGGACTTCCACATAATGG | 5' |
| HaloTag | /56-FAM/ACAGATCCT/ZEN/CAGTGGTTGGCTCGAG/3IABkFQ/ | 5' |
| MAPK8_fwd | CCCTCACTCCTCTGGGGTTATTTCA | 3' |
| MAPK8_rev | CTAACTGCAAGGCTGTGGACATCGG | 3' |
| HiBiT | /56-FAM/TGAGCGGCT/ZEN/GGCGGCTGTTCAAGAAGATTA/3IABkFQ/ | 3' |
| | **Chromosomal Probe for normalization** | |
| **Name** | **Sequence** | **Flank** |
| RPPH1_fwd | GTCATCAACCCGCTCCAAGGAATC | NA |
| RPPH1_rev | GACATGGGAGTGGAGTGACAGGAC | NA |
| RPPH1-Probe | /5Cy5/GGGCCCAGT/TAO/GTCACTAGGCGGGAA/3IAbRQSp/ | NA |
| | **dsDNA HDRT (TRAC/ GFP KI) synthesis** | |
| **Name** | **Sequence** | |
| TRAC-HDRT_fwd | TGAAATCATGGCCTCTTGGC | |
| TRAC-HDRT_rev | GGGAAGAAGGTGTCTTCTGGA | |

**Supplementary Table S3.** DNA donors used in this study.

| Type | Target | Insertion length (bp) | Sequence |
|---|---|---|---|
| ssDNA | gDel | 34 | A*A*CAGTTGGGCACTGACTTTATGCATAACATTTGCTAGATGTTGCTTAACTTCTCCTTTCGTTTAATTGAGTTGCAATTGGTTAATAACGGTATCCAAAGGACCCTCATTCTCGTCCTGATACTTTGGGTGTATCTGGATGTTTTTCACTTG*T*G |
| ssDNA | gIns | 34 | A*A*TGTTTCAGACCTTTCAACCACTAGCACTAGCCTTGCAACATCTCCCACTAAACCCTATGTTTAATTGAGTTGCAATTGGTTAATAACGGTATACATCATCTTCTCCTATCCTAAGTGACATCAAGGTGGGTGAATTGGGCCAAAAATGGC*A*G |
| ssDNA | gMej | 34 | T*T*TGTAGAAACATTTGAAAATGTTCCCTGGGTAGGTAACTCTGGGGTAGCAGTACCGTTGGTTTAATTGAGTTGCAATTGGTTAATAACGGTATTTGTCAAGACTCATGAACCCAGAAGCTATAGGGAAACGAGGAGGAAGAATCAGAACCT*A*A |
| dsDNA_fw | NA | 34 | G*T*TTAATTGAGTTGTCTAGAGTTAATAACGGT*A*T |
| dsDNA_rev | NA | 34 | A*T*ACCGTTATTAACTCTAGACAACTCAATTAA*A*C |
| ssDNA | HEK3 | 18 | G*T*CCTGCGACGCCCTCTGGAGGAAGCAGGGCTTCCTTTCCTCTGCCATCAATGATGGTGATGATGGTGCGTGCTCAGTCTGGGCCCCAAGGATTGACCCAGGCCAGGGCTGGAGAA*G*C |
| ssDNA | HEK4 | 6 | A*G*GGCCCCCACTGTAGTCACACAGCACCAGAGTCTCCGCTTTAACCCCCACATCTTCCTCCAGCCGCAGTGCCACCGGGGCGCCGCGGTGCCCCTGCCTGTCAT*C*C |
| ssDNA | HBEGF | 6 | G*T*GCAGGGTTCACAGGAAATGGAACCTGACCCATTAGGCATCAGCCCCCGCAATTGGTCAGGCAACATCACCCCTTCCCTGGGTAGGTGTGTGGGTGGAGGGGC*T*G |
| ssDNA-HiBiT | CTNNB1 | 39 | GGGCTGCCTCCAGGTGACAGCAATCAGCTGGCCTGGTTTGATACTGACCTGGTCTCCGTGAGCGGCTGGCGGCTGTTCAAGAAGATTAGCTAAATCATCGTTTAGGTAAGAAGTTTTAAAAAGCCAGTTTGGGTAAAATA |
| ssDNA-HiBiT | HDAC2 | 39 | CACATTGCCTCTTTTAACAGAACCAAATCAGAACAGCTCAGCAACCCCGTCTCCGTGAGCGGCTGGCGGCTGTTCAAGAAGATTAGCTGAATTTGACAGTCTCACGAATTTCAGAAAATCATTAAAAAGAAAATATTGAA |
| ssDNA-HiBiT | MAPK8 | 39 | CTGTCTGCAACTGATTTGCTGTTTTGTTTCTCATAGCACAGGTGCAGCAGGTCTCCGTGAGCGGCTGGCGGCTGTTCAAGAAGATTAGCTGATCAATGGCTCTCAGCATTCATCATCATCGTCGTCTGTCAATGATGTGTCT |
| ssDNA-HiBiT | NRAS | 39 | CAATTAACCCTGATTACTGGTTTCCAACAGGTTCTTGCTGGTGTGAAATGGTGAGCGGCTGGCGGCTGTTCAAGAAGATTAGCACTGAGTACAAACTAGTGGTGGTTGGAGCAGGTGGTGTTGGGAAAAGCGC |
| ssDNA-HiBiT | NR3C1 | 39 | TACCAAAATATTCAAATGGAAATATCAAAAAACTTCTGTTTCATCAAAAGGTCTCCGTGAGCGGCTGGCGGCTGTTCAAGAAGATTAGCTGACTGCGTTATTAAGAATCGTTGCCTTAAAGAAAGTCGAATTAATAGCTTTT |

| | | | |
|---|---|---|---|
| Plasmid-HaloTag-HiBiT | HDAC2 | 975 | CATAAGACAACATATGCCATCTTACCTATTGGTTTATCAGTATAAC<br>TATATCAGTCAGTTTGTATTCTTGTACGTAGTTTATCTGAGACCCT<br>TTTTTGCTGTTATCTTTTATATATGCACTTAATGGTTTTCATGGTG<br>TTCTTATTTCTGTGTACATGATTTTTTTAGTGTTGTGAGATAGGTA<br>AATTTAAGAAACAATTTTAAGTAAAAACAGTTGAGGATTTTAGTGT<br>CAGAAAGCATGTAAATGTTTGGGCACACATAATTTTGTGCCAGCCT<br>TTACAGGATAACTTGGAATAGTTTATTAAATCACTAGAAAAATTAA<br>GGCATTAATTACTCCAGAAACACTTAAGACATTATCAATTAATGTG<br>TTTCTTAGCATGAGCCTACAGGACCTGATCAATATAATAGGTCAGT<br>GCATTTAAACCAAGATTGTGCCATTTTTAATATTTCACATTGCCTC<br>TTTTAACAGAACCAAATCAGAACAGCTCAGCAACCCCCTCGAGCCA<br>ACCACTGAGGATCTGTACTTTCAGAGCGATAACGATGGATCCGAAA<br>TCGGTACTGGCTTTCCATTCGACCCCCATTATGTGGAAGTCCTGGG<br>CGAGCGCATGCACTACGTCGATGTTGGTCCGCGCGATGGCACCCCT<br>GTGCTGTTCCTGCACGGTAACCCGACCTCCTCCTACGTGTGGCGCA<br>ACATCATCCCGCATGTTGCACCGACCCATCGCTGCATTGCTCCAGA<br>CCTGATCGGTATGGGCAAATCCGACAAACCAGACCTGGGTTATTTC<br>TTCGACGACCACGTCCGCTTCATGGATGCCTTCATCGAAGCCCTGG<br>GTCTGGAAGAGGTCGTCCTGGTCATTCACGACTGGGGCTCCGCTCT<br>GGGTTTCCACTGGGCCAAGCGCAATCCAGAGCGCGTCAAAGGTATT<br>GCATTTATGGAGTTCATCCGCCCTATCCCGACCTGGGACGAATGGC<br>CAGAATTTGCCCGCGAGACCTTCCAGGCCTTCCGCACCACCGACGT<br>CGGCCGCAAGCTGATCATCGATCAGAACGTTTTTATCGAGGGTACG<br>CTGCCGATGGGTGTCGTCCGCCCGCTGACTGAAGTCGAGATGGACC<br>ATTACCGCGAGCCGTTCCTGAATCCTGTTGACCGCGAGCCACTGTG<br>GCGCTTCCCAAACGAGCTGCCAATCGCCGGTGAGCCAGCGAACATC<br>GTCGCGCTGGTCGAAGAATACATGGACTGGCTGCACCAGTCCCCTG<br>TCCCGAAGCTGCTGTTCTGGGGCACCCCAGGCGTTCTGATCCCACC<br>GGCCGAAGCCGCTCGCCTGGCCAAAAGCCTGCCTAACTGCAAGGCT<br>GTGGACATCGGCCCGGGTCTGAATCTGCTGCAAGAAGACAACCCGG<br>ACCTGATCGGCAGCGAGATCGCGCGCTGGCTGTCTACTCTGGAGAT<br>TTCCGGTGTCTCCGTGAGCGGCTGGCGGCTGTTCAAGAAGATTAGC<br>TGAATTTGACAGTCTCACCAATTTCAGAAAATCATTAAAAAGAAAA<br>TATTGAAAGGAAAATGTTTTCTTTTTGAAGACTTCTGGCTTCATTT<br>TATACTACTTTGGCATGGACTGTATTTATTTTCAAATGGCTTTTTC<br>GTTTTTGTTTTTCTTGGCAAGTTTTATTGTGAGTTTTTCTAATTAT<br>GAAGCAAAATTTCTTTTCTCCACCATGCTTTATGTGATAGTATTTA<br>AAATTGATGTGAGTTATTATGTCAAAAAAACTGATCTATTAAAGAA<br>GTAATTGGCCTTTCTGAGCTGATTTTTCCATCTTTTGTAATTATCT<br>TTATTAAAAAATTGTACTTGGATTATCTTTTGTCTGTTTATTACTA<br>CAATATGAAGTCTTGTTTCAGTGGCTAATGACATCATTTCTGTAGA<br>CTTACAATACACTCTAGGTGAAAGATAATGATTACAGCTTGAAAGA<br>TAACTATTTGCTGTTTCTTTGGGAAGAGTATTTATAGTAA |

| | | | |
|---|---|---|---|
| Plasmid-HaloTag-HiBiT | MAPK8 | 975 | TTGTTTCAAATTTGAATCATTTGAATTTTTCCTTTTTAGTGAGAAG<br>CTGTAAAGACTTTTTTGTAGGGAAGTAGCTTTTAACTTTTGTAGTT<br>ACACAGTCCTTTAAGATCCTCTGTCCAAAAAAAGGCATTACAGACA<br>GTTTTGCATGTATTATCAGCAGTATTCACACATACCCTGAAGCCCA<br>TTCATGGATCTTGCTGCAGGACCATTTCTAAATGTGGTTCAGATGT<br>AAAATTCTTGTCTTAAACTGAAAAACACATTCATTGAAAGGATAGG<br>ACTCCACGATTCTAGACATTTTCAGAATTCTCACCTCATAGCTGTC<br>AATGAAGAGTGTTTTTAAGTTAGTGTGTTGGATATCATTTGCGATT<br>ATTTTTAGTGAGCCTTCGAAACCCAAGAGAAAAAAATTACCACTGG<br>AGGCAGTCAGTGCAGTGCAAGTAGCTTGATCTGCAGCTGTCTGCAA<br>CTGATTTGCTGTTTTGTTTCTCATAGCACAGGTGCAGCAGCTCGAG<br>CCAACCACTGAGGATCTGTACTTTCAGAGCGATAACGATGGATCCG<br>AAATCGGTACTGGCTTTCCATTCGACCCCCATTATGTGGAAGTCCT<br>GGGCGAGCGCATGCACTACGTCGATGTTGGTCCGCGCGATGGCACC<br>CCTGTGCTGTTCCTGCACGGTAACCCGACCTCCTCCTACGTGTGGC<br>GCAACATCATCCCGCATGTTGCACCGACCCATCGCTGCATTGCTCC<br>AGACCTGATCGGTATGGGCAAATCCGACAAACCAGACCTGGGTTAT<br>TTCTTCGACGACCACGTCCGCTTCATGGATGCCTTCATCGAAGCCC<br>TGGGTCTGGAAGAGGTCGTCCTGGTCATTCACGACTGGGGCTCCGC<br>TCTGGGTTTCCACTGGGCCAAGCGCAATCCAGAGCGCGTCAAAGGT<br>ATTGCATTTATGGAGTTCATCCGCCCTATCCCGACCTGGGACGAAT<br>GGCCAGAATTTGCCCGCGAGACCTTCCAGGCCTTCCGCACCACCGA<br>CGTCGGCCGCAAGCTGATCATCGATCAGAACGTTTTTATCGAGGGT<br>ACGCTGCCGATGGGTGTCGTCCGCCCGCTGACTGAAGTCGAGATGG<br>ACCATTACCGCGAGCCGTTCCTGAATCCTGTTGACCGCGAGCCACT<br>GTGGCGCTTCCCAAACGAGCTGCCAATCGCCGGTGAGCCAGCGAAC<br>ATCGTCGCGCTGGTCGAAGAATACATGGACTGGCTGCACCAGTCCC<br>CTGTCCCGAAGCTGCTGTTCTGGGGCACCCCAGGCGTTCTGATCCC<br>ACCGGCCGAAGCCGCTCGCCTGGCCAAAAGCCTGCCTAACTGCAAG<br>GCTGTGGACATCGGCCCGGGTCTGAATCTGCTGCAAGAAGACAACC<br>CGGACCTGATCGGCAGCGAGATCGCGCGCTGGCTGTCTACTCTGGA<br>GATTTCCGGTGTCTCCGTGAGCGGCTGGCGGCTGTTCAAGAAGATT<br>AGCTGATCAATGGCTCTCAGCATCGATCATCATCGTCGTCTGTCAA<br>TGATGTGTCTTCAATGTCAACAGATCCGACTTTGGCCTCTGATACA<br>GACAGCAGTCTAGAAGCAGCAGCTGGGCCTCTGGGCTGCTGTAGAT<br>GACTACTTGGGCCATCGGGGGGTGGGAGGGATGGGGAGTCGGTTAG<br>TCATTGATAGAACTACTTTGAAAACAATTCAGTGGTCTTATTTTTG<br>GGTGATTTTTCAAAAAATGTAGAATTCATTTTGTAGTAAAGTAGTT<br>TATTTTTTTTAATTTCAAGTGATGTAATTTAAAACCTAAGTTGTGT<br>TTCAAAACAGCAACAAAACTGTATTGTATTTTTTTTGCTGTAATTA<br>ACTGTATAATGTAAACCTAATTATTTTATCATGGTTTAAATTTTTT<br>GCATATTTGCTTTATCTTATGCTGCTGATTTTTTTAACTGAATTTG<br>TAAGATTTTGTTTATCAAAGCAACTATTATGTGGTGACTTGCC |

| | | | |
|---|---|---|---|
| Plasmid-HaloTag-HiBiT | NRAS | 975 | GAGGTAAGTTTATCTCATGCATAGTGTTCGGCTTTGGGCTGTGGAA<br>TGTTCAGGCGTTTCACTGATGCCAGAAATGGAGCAGAATCTATCAG<br>CTGGAGACAAAGGCCTTGGGCGGGGGTCCTTCCATTTGGTGCCTAC<br>GTGGGGAGATCTTTGGAGACAGAAGGGAGAATGGGAAGGAGTTGCG<br>GCCTGGAGGCTTCCTGCTAGAGCTGAGAAGCCTTCGGGGAGTAATA<br>GGAAGGGGGATTTCCATTGCTTAGGCTGAGGGCGGGGCCCAAGGAC<br>TGTTGAAAAATAGCTAAGGATGGGGGTTGCTAGAAAACTACTCCAG<br>AAGTGTGAGGCCGATATTAATCCGGTGTTTTTGCGTTCTCTAGTCA<br>CTTTAAGAACCAAATGGAAGGTCACACTAGGGTTTTCATTTCCATT<br>GATTATAGAAAGCTTTAAAGTACTGTAGATGTGGCTCGCCAATTAA<br>CCCTGATTACTGGTTTCCAACAGGTTCTTGCTGGTGTGAAATGGTG<br>AGCGGCTGGCGGCTGTTCAAGAAGATTAGCGCAGAAATCGGTACTG<br>GCTTTCCATTCGACCCCCATTATGTGGAAGTCCTGGGCGAGCGCAT<br>GCACTACGTCGATGTTGGTCCGCGCGATGGCACCCCTGTGCTGTTC<br>CTGCACGGTAACCCGACCTCCTCCTACGTGTGGCGCAACATCATCC<br>CGCATGTTGCACCGACCCATCGCTGCATTGCTCCAGACCTGATCGG<br>TATGGGCAAATCCGACAAACCAGACCTGGGTTATTTCTTCGACGAC<br>CACGTCCGCTTCATGGATGCCTTCATCGAAGCCCTGGGTCTGGAAG<br>AGGTCGTCCTGGTCATTCACGACTGGGGCTCCGCTCTGGGTTTCCA<br>CTGGGCCAAGCGCAATCCAGAGCGCGTCAAAGGTATTGCATTTATG<br>GAGTTCATCCGCCCTATCCCGACCTGGGACGAATGGCCAGAATTTG<br>CCCGCGAGACCTTCCAGGCCTTCCGCACCACCGACGTCGGCCGCAA<br>GCTGATCATCGATCAGAACGTTTTTATCGAGGGTACGCTGCCGATG<br>GGTGTCGTCCGCCCGCTGACTGAAGTCGAGATGGACCATTACCGCG<br>AGCCGTTCCTGAATCCTGTTGACCGCGAGCCACTGTGGCGCTTCCC<br>AAACGAGCTGCCAATCGCCGGTGAGCCAGCGAACATCGTCGCGCTG<br>GTCGAAGAATACATGGACTGGCTGCACCAGTCCCCTGTCCCGAAGC<br>TGCTGTTCTGGGGCACCCCAGGCGTTCTGATCCCACCGGCCGAAGC<br>CGCTCGCCTGGCCAAAAGCCTGCCTAACTGCAAGGCTGTGGACATC<br>GGCCCGGGTCTGAATCTGCTGCAAGAAGACAACCCGGACCTGATCG<br>GCAGCGAGATCGCGCGCTGGCTGTCGACGCTCGAGATTTCCGGCGA<br>GCCAACCACTGAGGATCTGTACTTTCAGAGCGATAACACTGAGTAC<br>AAACTGGTGGTGGTTGGAGCAGGTGGTGTTGGGAAAAGCGCACTGA<br>CAATCCAGCTAATCCAGAACCACTTTGTAGATGAATATGATCCCAC<br>CATAGAGGTGAGGCCCAGTGGTAGCCCGCTGACCTGATCCTGTCTC<br>TCACTTGTCGGATCATCTTTACCCATATTCTGTATTAAAGGAATAA<br>GAGGAGAGAAAGTAAAAAGTTATTTTGGGTATACATTCAGTTATGC<br>AATAAGCTTAACGTGTTTATAGAGAACAGTTCATTTTTATTAGCTG<br>CTGAAGTTTCTAAAACCTGTCCAGTTTTTAACAGTTCTGTAAACTA<br>TTGCAAACTCAGTGTTGAGTTCATTCATGAGTTTCTTCATATATAA<br>CAGCTCTATTACATGAGAAACACAGGCCATAGTAGCGAGACTGTCT<br>GATTGTATGGGAGATAATAGGATGGAGATAAAGGATTCAGAGATGA<br>GTGTTCTTCAATATTTATTTATTAGCTAGTT |

| | | | |
|---|---|---|---|
| Plasmid-HaloTag-HiBiT | NR3C1 | 975 | ATTATGAAATATCCTTTCACCTAGTCATGTGTATATAAAATCACCA<br>TGTTATTACAGAATTTAGTAATACTGTTTTTAAAAAGTATGATTAA<br>TCCATTAAATTAGAATAATGCACCCTTCATATATTATGGTACTACA<br>GTGATTCATGAAATAATTCTATATAATTCTACATACAATCAAAGAA<br>ATATAAAATGTGTTTTGTACGGAAGTGCTTATTTTTCATCTGGGGA<br>ATTCCAGTGAGATTGGTATATTCTAGGCCAGATAATTTTTTCAAAA<br>TAGAGGACAACAAACATGAGATGTTCCCACTGACCAATTTGGAAGC<br>CTGATCATTACCATATCTTCTCTTGCAGGTGGTTGAAAATCTCCTT<br>AACTATTGCTTCCAAACATTTTTGGATAAGACCATGAGTATTGAAT<br>TCCCCGAGATGTTAGCTGAAATCATCACCAATCAGATACCAAAATA<br>TTCAAATGGAAATATCAAAAAACTTCTGTTTCATCAAAAGCTCGAG<br>CCAACCACTGAGGATCTGTACTTTCAGAGCGATAACGATGGATCCG<br>AAATCGGTACTGGCTTTCCATTCGACCCCCATTATGTGGAAGTCCT<br>GGGCGAGCGCATGCACTACGTCGATGTTGGTCCGCGCGATGGCACC<br>CCTGTGCTGTTCCTGCACGGTAACCCGACCTCCTCCTACGTGTGGC<br>GCAACATCATCCCGCATGTTGCACCGACCCATCGCTGCATTGCTCC<br>AGACCTGATCGGTATGGGCAAATCCGACAAACCAGACCTGGGTTAT<br>TTCTTCGACGACCACGTCCGCTTCATGGATGCCTTCATCGAAGCCC<br>TGGGTCTGGAAGAGGTCGTCCTGGTCATTCACGACTGGGGCTCCGC<br>TCTGGGTTTCCACTGGGCCAAGCGCAATCCAGAGCGCGTCAAAGGT<br>ATTGCATTTATGGAGTTCATCCGCCCTATCCCGACCTGGGACGAAT<br>GGCCAGAATTTGCCCGCGAGACCTTCCAGGCCTTCCGCACCACCGA<br>CGTCGGCCGCAAGCTGATCATCGATCAGAACGTTTTTATCGAGGGT<br>ACGCTGCCGATGGGTGTCGTCCGCCCGCTGACTGAAGTCGAGATGG<br>ACCATTACCGCGAGCCGTTCCTGAATCCTGTTGACCGCGAGCCACT<br>GTGGCGCTTCCCAAACGAGCTGCCAATCGCCGGTGAGCCAGCGAAC<br>ATCGTCGCGCTGGTCGAAGAATACATGGACTGGCTGCACCAGTCCC<br>CTGTCCCGAAGCTGCTGTTCTGGGGCACCCCAGGCGTTCTGATCCC<br>ACCGGCCGAAGCCGCTCGCCTGGCCAAAAGCCTGCCTAACTGCAAG<br>GCTGTGGACATCGGCCCGGGTCTGAATCTGCTGCAAGAAGACAACC<br>CGGACCTGATCGGCAGCGAGATCGCGCGCTGGCTGTCTACTCTGGA<br>GATTTCCGGTGTCTCCGTGAGCGGCTGGCGGCTGTTCAAGAAGATT<br>AGCTGACTGCGTTAATAAGAATGGTTGCCTTAAAGAAAGTCGAATT<br>AATAGCTTTTATTGTATAAACTATCAGTTTGTCCTGTAGAGGTTTT<br>GTTGTTTTATTTTTTATTGTTTTCATCTGTTGTTTTGTTTTAAATA<br>CGCACTACATGTGGTTTATAGAGGGCCAAGACTTGGCAACAGAAGC<br>AGTTGAGTCGTCATCACTTTTCAGTGATGGGAGAGTAGATGGTGAA<br>ATTTATTAGTTAATATATCCCAGAAATTAGAAACCTTAATATGTGG<br>ACGTAATCTCCACAGTCAAAGAAGGATGGCACCTAAACCACCAGTG<br>CCCAAAGTCTGTGTGATGAACTTTCTCTTCATACTTTTTTTCACAG<br>TTGGCTGGATGAAATTTTCTAGACTTTCTGTTGGTGTATCCCCCCC<br>CTGTATAGTTAGGATAGCATTTTTGATTTATGCATGGAAACCTGAA<br>AAAAAGTTTACAAGTGTATATCAGAAAAGGGAAGTTGTGCCTT |
| Electroporation enhancer oligo | | | CCACCAGGACACCCCCATCCCCCACCCCCCCCTCCTCCTCCCGGAC<br>AACCACTACCTCACCACCCACTCCCCCCTCACCAAACACCCCAAGG<br>ACA |

**Supplementary Table S4.** Reference sequences used in this study.

| Target | Reference sequence |
|---|---|
| gMej | CCTGAAGTCAGGAAATTGCATCAGGTGATATCACCAGGCTGATATCTTTAGGTTCTGATT<br>CTTCCTCCTCGTTTCCCTATAGCTTCTGGGTTCATGAGTCTTGACAACAACGGTACTGCT<br>ACCCCAGAGTTACCTACCCAGGGAACATTTTCAAATGTTTCTACAAATGTATCCTACCAA<br>GAAACTACAACACCTAGTACCCTTGGAAGTACCAGCCT |
| gIns | TGAGGGGCAATCTGCCATTTTTGGCCCAATTCACCCACCTTGATGTCACTTAGGATAGGA<br>GAAGATGATGTATAGGGTTTAGTGGGAGATGTTGCAAGGCTAGTGCTAGTGGTTGAAAGG<br>TCTGAAACATTTCCAGGTGACAGGCTAGGCTTCAAGGTTGTCTCTGGAGTTGAAACGTTG<br>GCTGG |
| gDel | GGCATTCTCTGAAGAGTGGGTTTCACAAGTGAAAAACATCCAGATACACCCAAAGTATCA<br>GGACGAGAATGAGGGTCCTTTGGGAAAGGAGAAGTTAAGCAACATCTAGCAAATGTTATG<br>CATAAAGTCAGTGCCCAACTGTTATAGGTTGTTGGATAAATCAGTGGTTATTTAGGGAAC<br>TGCTTGACGTAGGAA |
| HBEGF | GCAACATTATGCCTGCCAGAGAGCATGGCAGTGCTGTCATCCCTGAGGGGTTGCCAAGGC<br>AGCCGTTTGCTGGGTTTGGCTTAGGAGAGAGGCTAAGGAACCCACAGCCCCTCCACCCAC<br>ACACCTACCCAGGGAAGGGGTGATGTTGCCTGACCGGGGGCTGATGCCTAATGGGTCAGG<br>TTCCATTTCCTGTGAACCCTGCACAAAGGAATTGTTCTCTCACAACGTGGCC |
| PCSK9 | TGGTATTCATCCGCCCGGTACCGTGGAGGGGTAATCCGCTCCAGGTTCCACGGGATGCTC<br>TGGGCAAAGACAGAGGAGTCCTCCTCGATGTAGTCGACATGGGGCAACTTCAAGGCCTGC<br>AGAAGCCAGAGAGGCCGGGGGACTCTGCTTAAGCCATTTGGACAGCAGCAAACACAGCCA<br>CAGCCTGGCCTTACCTGATCAAACCTGTCCCCACAT |
| HEK3_off1 | GGAGAAGCATGAACCAGTCAAAAAGTTTAAAGACAAGAGCATTAACTGCACCAGTGGGCA<br>GCTCAGCTCAGACACCAGTAGCGTGGGCACCCAGACTGAGCACGTGCTGGAGCCCAAGAA<br>ATGCAGAGACCTGTGCACCTCTGGTCAGGGCAAGTACAGTGACAGGCACACCATGAAGCA<br>CTCAGACGATGACTCAGAAATTGTCAGCGACGACATCAGTGACATTTTCCGATTTCTTGA<br>TGACATGAGCATCAGTGGCTCCACG |
| HEK3_off2 | TGGGAGAGTTAACCTGGGTTCAGTAGACAGGGCCCAGGACCACAGTGAACTAAAGCATAG<br>TTCAGGAGTACACAGAGGTCCAAGGAGGCCTATGCAGGGGACAGAACACTGATGAGTAGC<br>AGATGACACTTGGTGTTGACAGGGAGCAACTTCACAGTCCCAGGCATCAGGACACAGACC<br>GGGCACGTGAGGGAAGCCCAAGGGAGAGGACTGGTGTAATCGAGGCTGACTCCACTTTTA<br>ATGTTTGACTGATGATAGGTTTCAAGTCTCACTAAGTCTC |
| HEK3_off3 | CAGCCGGCGATGAGTAAGAGTGATGTGGGAGGTTCCTGAGGATGCGGGTGCTGGGAAGAG<br>GGCTGGCCCAGATTTCCAGGGACTGAGAGGGAACAGAAGGGCTAAGACTAAAAGGAACAG<br>AGGAGTTCATAGTGAGCGGTAAAGAGCTCAGACTGAGCAAGTGAGGGGCTCAGCCTCCCA<br>TGGAGGACAGGGGGCTGGGGCCCCTGGCTGATGTCTGGACTGAAGCCCCCACGCCCAGAG<br>GTTCTTGGGCCTTTGGACCAGGAAT |
| HEK3_off4 | ACAAAAGCTATGATGTGATGTGACTGGGCATGGTGTCTCACCCCTGTAATCCTAGCACTT<br>TGGAAGGTCGAAGCGGCAGGATGGCTTCAACCCAGGAGTTCGAGACCAGACTGAGCAAGA<br>GAGGGAGAGTGTCTGTATTAACAACAAACAAACAAACAAAAAACTAAACTAAAAGAAACT<br>GTGGTGTATAATATAAAATTCTGGCTGAGCAGATTAAGATGAG |
| HEK3_on | CCATGCAATTAGTCTATTTCTGCTGCAAGTAAGCATGCATTTGTAGGCTTGATGCTTTTT<br>TTCTGCTTCTCCAGCCCTGGCCTGGGTCAATCCTTGGGGCCCAGACTGAGCACGTGATGG<br>CAGAGGAAAGGAAGCCCTGCTTCCTCCAGAGGGCGTCGCAGGACAGCTTTTCCTAGACAG<br>GGGCTAGTATGTGCAGCTCCTGCACCGGGATACTGGTTGACAAGTTTGGCTGGGCTG |
| HEK4_off1 | CTGGGGCTGAAGATCCCTAGGGGGGCTCTGCTGGGCTCACTGCTCTCCAGAGTGGTCCAG<br>CCCGGCTGCAGGGTGCTGCTTCCAGCTTGGTGCACTGCGGCCGGAGGAGGTGGAGGATGG<br>AAAGTAAGATTCAAAGACAGGGAGTGCAAGGGGACAGTGATACTTGAGGACTCCGAGGAG<br>GAAGTAGGAGAATCATCCGGAATG |
| HEK4_off2 | GTGTGTGTGTTTGTGTGTGTGCCTGGGGCAACCAACATGGTGGGACACTCCCTGCAGGCT<br>GTGGTGAAGAGGATGGGGTGATCCCCAGGCCCTGGGTGGTGTGCTTGGGTTGCTTTGGCA<br>ATGGAGGCATTGGGCAGGGGAAGCCTGTCTTCAGGGCACATGCACGTGCGCAGGGCTCTG<br>CGGCTGGAGGGGGTGGGGTTGCTGTTAGTGACAGGGGCCCCAGCCAGGCAGGTTTCAGGA<br>TTGGGGAGCACTTGCTTCGGCTCCCTTGCTCTCATG |

| | |
|---|---|
| HEK4_off3 | GTGAGCTCATTTCCACCAGAACTCAGCCCAGGCTGCTGTGGGATGGAATCACCTGCACCC<br>GGATGTTCTTTCTGGGCTGGTACATACAGGCAAGGCATCACGGCTGGAGGTGGAGGGGGC<br>CTAACCCGGGGTTGCCCAGGAAGGGGTTTGCACATGGATTCGGTGTGTTGTGGAGGAACC<br>GAGGGTGGAAACAGCTGGGCCACCCCACACCTAAGCCCGTATTTCCCATCTGTCCTTATC<br>TACTCTGC |
| HEK4_on | TGGAAGGAAGGGAGGAAGGGCGAGGCAGAGGGTCCAAAGCAGGATGACAGGCAGGGGCAC<br>CGCGGCGCCCCGGTGGCACTGCGGCTGGAGGTGGGGGTTAAAGCGGAGACTCTGGTGCTG<br>TGTGACTACAGTGGGGGCCCTGCCCTCTCTGAGCCCCCGCCTCCAGGCCTGTGTGTGTGT<br>CTCCGTTCGGGTTGAAAGG |
| CTNNB1 | CCTAAACCACTCCCACCCTACCAACCAAGTCTTTCTGAAGTTCTGTAGGCAGAGTAAAAG<br>TATTTTACCCAAACTGGCTTTTTAAAACTTCTTACCTAAAGGATGATTTACAGGTCAGTA<br>TCAAACCAGGCCAGCTGATTGCTGTCACCTGGAGGCAGCCCATCCATGAGGTCCTGGGCA<br>TGCCCCAGATCTGGCAGCCCATCAACTGGATAGTCAGCACCAGGGTGGTGGCCACCCATC<br>TCATGTTCCATCATGGGGTCCATACCCAAGGCATCCTGGCCATATCCACCAGAGTGAA |
| HDAC2 | CCATGCCAAAGTAGTATAAAATGAAGCCAGAAGTCTTCAAAAAGAAAACATTTTCCTTTC<br>AATATTTTCTTTTTAATGATTTTCTGAAATTGGTGAGACTGTCAAATTCAGGGGTTGCTG<br>AGCTGTTCTGATTTGGTTCTGTTAAAAGAGGCAATGTGAAATATTAAAAATGGCACAATC<br>TTGGTTTAAATGCACTGACCTATTATATTGATCAGGTCCTGTAGGCTCATGCTAAGAAAC<br>ACATTAATTGATAATGTCTTAAGTGTTTCTGGAGTAATTAATGCCTTAATTTTTCTAGTG<br>ATTTAATAAACTATTCCAAGTTATCCTGTAAAGGCTGGCACA |
| MAPK8 | AATGACTAACCGACTCCCCATCCCTCCCACCCCCCGATGGCCCAAGTAGTCATCTACAGC<br>AGCCCAGAGGCCCAGCTGCTGCTTCTAGACTGCTGTCTGTATCAGAGGCCAAAGTCGGAT<br>CTGTTGACATTGAAGACACATCATTGACAGACGACGATGATGATGGATGCTGAGAGCCAT<br>TGATCACTGCTGCACCTGTGCTATGAGAAACAAAACAGCAAATCAGTTGCAGACAGCTGC<br>AGATCAAGCTACTTGCACTGCACTGACTGCC |
| NRAS | ACTACTCCAGAAGTGTGAGGCCGATATTAATCCGGTGTTTTTGCGTTCTCTAGTCACTTT<br>AAGAACCAAATGGAAGGTCACACTAGGGTTTTCATTTCCATTGATTATAGAAAGCTTTAA<br>AGTACTGTAGATGTGGCTCGCCAATTAACCCTGATTACTGGTTTCCAACAGGTTCTTGCT<br>GGTGTGAAATGACTGAGTACAAACTGGTGGTGGTTGGAGCAGGTGGTGTTGGGAAAAGCG<br>CACTGACAATCCAGCTAATCCAGAACCACTTTGTAGATGAATATGATCCCACCATAGAGG<br>TGAGGCCCAGTGGTAGCCCGCTGACCTGATCCTGTCTCTCACTTGTCGG |
| NR3C1 | TGACGACTCAACTGCTTCTGTTGCCAAGTCTTGGCCCTCTATAAACCACATGTAGTGCGT<br>ATTTAAAACAAAACAACAGATGAAAACAATAAAAAATAAAACAACAAAACCTCTACAGGA<br>CAAACTGATAGTTTATACAATAAAAGCTATTAATTCGACTTTCTTTAAGGCAACCATTCT<br>TATTAAGGCAGTCACTTTTGATGAAACAGAAGTTTTTTGATATTTCCATTTGAATATTTT<br>GGTATCTGATTGGTGATGATTTCAGCTAACATCTCGGGGAATTCAATACTCATGGTCTTA<br>TCCAAAAATGTTTGGAAGCAATAGTTAAGGAGATTTTCAACCACCTGCAAGAGAAGATAT<br>GGTAATGATCAGGCTTCCAAATTGGTCAGTGGG |
| HBEGF long-read | TTGCCCATGACCTGAACAGCCTGGATTCTCCTGGCCCTCTCCTCCTAGGCTGGGCAGGGC<br>TGGGCTGTGACTCACCCCACCCCCACCCCCCACCCACACGGCTGCTCCTCTTACCTCTGC<br>AGACCTGACTCACTGCTCCCTGTCCATGGCAGGAGCCTGGCTGTCACCCTGCACCTTCTC<br>CCTCCCCTTTCTGATTGGCTTGGCCCCCCTGCCTTGCTCTCCCCGAAGCTCTGGTCACTG<br>GGTTCCTCTGACCACCTGTATCACCTTCTGAGCTCTGAGGGGGCCTGGGACTGGATGAGA<br>GGAAATGAAAGACTGTGGGGGCTGCTGGCACCTACTTCTCTTCCCTTCTTTTGGCTTTGC<br>TGGGCAAGGACTATTTTTCAGGTCTGGGGATCCTACCACCTAAAATAAATGACTGCTACC<br>ATTTATTAAATTCCTACTGTGTTCTAGGCACTTGATATGTTATCCTGGCTAATGTAACAC<br>TTATAGCAACCTTTTGAGATAGTTACTTTGGCTATCCACATTTTACTGAGAACCTGAGGT<br>TCAGAGGAGTTAAGTGACTGCCCACAGTAAATAGCTGAAATTGGAGCACAGGTCTATGGA<br>CTTCAGAGCCCATTCATGCCTGGATCAGCATCTCAGGTGCTCTAGACTTGTGAGAGGGAG<br>GAGATGGGAGTGTGTGAGGCAGCTTGGTGTGGTGAGGAAGGACATTGGAGTGAAGTCCAG<br>AGAACACAGTTCTAATCCCAATCCTGCATGACCTTGAGTAAGTCACTCTGCCTGCCATGA<br>GTTTTTTCTTTTTTTCTTTTTTTTTTTTTTAAACATAGTCTCACTCTGTCACCCAGGCTG<br>GAGTGCAATGGCACGATCTCAGCTCACTGCAATCTCTGCCTCCCAGGTTCAAGTGATTCT<br>CCTGCCTCAGCCTCCTGAGTAGCTGCGATAACAGGCACACACCACCACGCCCGGCTAATT<br>TTTGTATTTTTTGTAGAGATGAGATTTTTGCCATGTTGGCAAGGCTGGTCTCGAACTCCC<br>GACCTCAGGTGATCCACCTGCCTCAGCCTCCCAAAGTGTTGGGATTACAGGCGTGAGCCA<br>CCGTGCCTGGCCACATGGTATTCTTTGAAGTCCCTCTAGCTTGAGACTCTAAGTCTCTAG<br>TCTAACGTATCATGCTTACCCTTCTGTAAGACACATGGCTGTAGCCATGGATGTGGGCAC<br>CTTTTTCCTGATGGGGGATAAAAGGGTGGATTGGGCTGATAGGCATAGTCCCTGGTCAA<br>TCCCAGCTGGATATCTGGGTGAGGCTGTTTTTCCCCCAGTCTCTCTGAAGCATGGAAAGA |

AGGAGGGAGTCATCATTGTTCCAGTTCCTTCTGGACAGTTCCTTACTTTCCATTTTTCTA
TCCCTTGTACACCCTGTACCCCCCAATCCAGAGAGCTATAAACAGGACATTGGGGGTTAA
ATATGAATGAATCTTTGAGAAAGTGGGTGAGCTGTAAAGGGTATGCAAGTTAAATATTTT
GCTTGAAGTTGAAAAAGCAAGGCCGTGACCAGGGCTGGCCTGCTTGCTGTTCCTGAGCCA
GGCTCTGCCCTGGGCTCATAGTACTAAGGGGTGCCCCAGAAGAGACCACCTGAACACATG
GACACTGTTCTTATATTAGGAGCCCTCCAACCCCAGAACCTCCAAGTACCTTCTCTAGAA
GCAATTTTTGTGTGTGACACTGTCTTTCTGCAAGTGGTTCACTGAGTACAGCATCAGGAA
ATGAGGCTGATTGAAGGCCAAAATAGAATGAAGTGGGTGTGGGGGAGTAGGAGATGGGGG
TGTAAGGTGGACAGTGGGGTGGAGGTGAGGTTGGTAGAATTGCCCAGTTACTCAACAAAA
GCATTCTGAGAATGAGGCTCTTACACAGAGACTGTGAAATGCCTTCCTTGGGACCCACCC
TAGCTTCTACTTCCTACCGAGGTTCCCTCTTTCTGGTGGTTCTGCCCAATCTTCCTGCTC
TTCCTTCTGCCTCTTAGGAGGCACTGAGCTAAGGGGCCTTCCCAGATCTCTGACTTCAGG
TGGAATCAAAGCATATATACTCCTTTCAAGCACTATGCTCTTCTGATTTTCTTCCCAAAG
AGTCAGACTTTAACAGAGTGCTTTTCTCCTACAGTCACTTTATCCTCCAAGCCACAAGCA
CTGGCCACACCAAACAAGGAGGAGCACGGGAAAAGAAAGAAGAAAGGCAAGGGGCTAGGG
AAGAAGAGGGACCCATGTCTTCGGAAATACAAGGACTTCTGCATCCATGGAGAATGCAAA
TATGTGAAGGAGCTCCGGGCTCCCTCCTGCATGTAAGTGCCCCTTCCCCAGGGCTGAATC
TCATCAGCACACTTTGTCAGCCACGTGGCTGTTCCTCGTTGTCACTGTTCCTTGAATTCA
TAATTTCACCCAGTTTCTTCTCAACCTCTGGGCGGAAGTTGGGAGGAGGGGAAATATATT
TTTAGTCAGCGGAAGCCCCCTCCCCCCTATAGGATGCAATTTCCTGTGGTATGGTTTTGT
GACGTGCTTTAATCCTTGGGGACATTTCCTGCTTGCCCAGAAATGAGCATGTGGCTAGGA
CAGCTGGCACCTGAAGGCAGGCCCTTAATTCTTGCCTGATGCCCTACTCTGGGAGGGAGA
AGCCAGTAGGAAACATGGCAGAGTGGGCTTCCAGGGCAGAGTAGAGCTCCTGTGGGAAGG
TAGGAAGTGCATTTGGATGCATGATGTATAGGTATGTGTGTATTTGGGTTTATGTGCATG
TAAGTGTGCAAATGTGGATTGACTGTGAGGCATGGCAGGACTGTACAGAGAGGGATCATC
ATGGCGGCAGGTTGAGGCCTCTCTTTCTTCTTCCTTATCCCAGCAAGGACGAGGAGGTGG
GAGACATGGAGAGTACTGGCCTTTGGCCACGTTGTGAGAGAACAATTCCTTTGTGCAGGG
TTCACAGGAAATGGAACCTGACCCATTAGGCATCAGCCCCCGGTCAGGCAACATCACCCC
TTCCCTGGGTAGGTGTGTGGGTGGAGGGGCTGTGGGTTCCTTAGCCTCTCTCCTAAGCCA
AACCCAGCAAACGGCTGCCTTGGCAACCCCTCAGGGATGACAGCACTGCCATGCTCTCTG
GCAGGCATAATGTTGCCACTGTGCCTGAGGCCAACACCCTGCGTCAGGCTGCAAACATCC
ATTCCCTTCCCTGTGGGGAGGGAGGCTCTGGGGGCCTTAGTGGGAGACTCTGGACAGGGC
CAAGAGACTGTTGTATGCACACTGCCTCCAGCCTGTCAAGAAGGCGGCGTGCCTGGCATC
CCTTCTACTGGTGATTGGTGCAGATCCCTTAGCTTTTTAAAGCTTCCTTGTTTTGTCTGA
TCACACACAGCAGAGCTGCCCTGTATTTGGCAGTTGGCAGACAGACCCATCACTCCCCAC
CATGTCCACAGTCACTTGTGCATCCTTTCCTATAACATCCTTGTCAGGAGCTTGGTATTA
GAGGGAGTTGTTTAAGAGTGGCATAGAAAGCCCCCATATTATCCTTCCCAAGGTCTTGGG
ACAGGGTGGGAAATGTTCATCTTAAATTTGTAAAATGGCATCATTAGTACAGGGTGAAGA
AGGTGACTCAAGTAGTCAAGGTGGATTGAGGTCAGGAATCTGTCTATACCAGATTGGTCC
TGGGCATTTTGGTGGATGGATGTGGGGCTTGCACTGTGTGGTTGAGAGGCCTTATAAGGT
TGCCCTCCTGGAGAGCTGGACTCGGATGACCACCTAAACCCAGAGAACCTGATATGGGTG
CCCAGGCCACCTTCCCAGTGGTCCCTAGGGATAGTGATAACTATAATGATGTCATATCTC
CTTTGTCCCAGAGTTTCAGTGTTTATATATAATATGAGTTGAGCCCAAGTATGTTGAGCC
CCTATTTGGTGGCAGACACTACTTTAGGAGCTGGAGAGATATAGTTTCCTGGGATTTTTC
AAAAGCCCTCTGCTGAGTAGGCAGGACTTGGTACCTCTACTTGAAAGGTGATGAAACTGG
AGCCAGAAAATAGGAAGTAATTTGCCTGAGGTCAATAGCTAAATAAGTAGTTGGAAATAA
GACAGAGTCTCAGTACCTGACTCCTAGTCCAACATGCTTTTCATGCCCTCAAGCTGTACT
GGGTGTTGGCTTTCATCTTTCTTTCCTGTATCTGTCCTTATAGAGTTGGAGCAGCATTTT
ATAGAGGGCAGAGGGCAGCTGTTGTCCTAGAGGTCTCTTATTCTTTTACTAGTCTAACAG
CACAGCAATCTGATTTGAAAACTTTACATTAACTTCTTGGGCAGAATTTTCTTTTTCTTT
GTTCTTTTCTTTCTTTCTTTCTTTTTTTTTTTTTTTTTTTTTTTTTGAGACAGAGTCTCACT
CTGTCTCCCATGCTGGGGTGCAGTGGTGTGATCTCAGCTCACTGCAACCTCTGCCTCCTG
GGTTCAAGCAATTCTCCTGCCTCAGCCTCCTAAGTGGCTGGGACTACAGGCACCTGCCAC
CATGCCGAATTAATAATTTTTATATTTTTAGTAGAGACGTAGTTTTGCCGTGTTGGCCAG
GCTGGTCTTGAACTCTTGACCTCAGGTGATCCGCCTGCCTCAGCCTCCCAAAGTGCTGGG
ATTACAGGCATGAGCCACCATATCTAGCCTTTTTTTTTTTTGAGATGGAATCTCGCTCTG
TCACCCAGGCTGGAGTGCAGTGACACAATCTCGGCTCTCTGCAGCCTCCGCCTCCCAGAT
TAAAGTGATTTTCCTGCTTCAGCCTCCTGAGCAGCTGGTATTACAGGCACATGCCCCCAC
ATCTGGCTAATTTTTAAATTTTTGTGGAGATGGGGTTTCACCATGTTGGCCAGGCTGGTC
TTGAACTCCTAACCTCAAGTAATCAGCCTGCCTTGGACTCCCAAAGTGCTGGGATTACAG
GCGTGGGCCACCACTTCCTGGGCAGATTTTCAGGGGGTTGATTGCATGTCTGGACTGGCC

| | |
|---|---|
| | CCCTACTGCCTCCTGCCCTTGCTACTCAGGGCAGAAAGCAGCAAGAAGACAGAAATCCTG<br>GTTTGGGGGAATGTGACATCTGTGCACGTTCATCTGGGGATCTTTGTGGCTCTTGTTTGA<br>CTCCAGACCCAGGAACCACTAGCCAGGGTGTGTCCAGGCTGCTGTGGTGAGCCTGAGGCT<br>AGCTGGCTTCCTAAACTAGCCCTCTGCAGCCACCATGAACAGGAAAACCCTTTTTGTGTC<br>ACCAGCCAAAAGTTGCCCTCAAAGAGTAGTTTCTGCTGGGCACAGTGGCTCACACCTGTA<br>ATCACAGCACTTTGGGAGGCCGAGGCACGTGGGTCGCCTGAGGTCAGGAGTTCGAGACCA<br>GCCTGGCCAACATAGAGAAACCCCCGTCTCTACTAAAAATACAAAAATTAGCTGGGTGTT<br>GTGGCGGGCGCCTGTAATCTCAGCTACTAGAGAGGCTGAGGCAGGAGAATCTCTCAAACC<br>CAGGAGGCAGAACTTGCAGTGAGCCGAGATAGTGCCATTGCACTCCAGCCTAGGCAACAA<br>GAGCAAAACTCCATCTCAAAAAAATAATAATAATAAATAAATAAAAGAGTAGTTTCCTGG<br>GATTCCTGACTAGTTGCCTACCCAGAAATTGGCT |
